# Versatile, do-it-yourself, low-cost spinning disk confocal microscope

**DOI:** 10.1101/2021.09.04.458950

**Authors:** Aaron R. Halpern, Min Yen Lee, Marco D. Howard, Marcus A. Woodworth, Philip R. Nicovich, Joshua C. Vaughan

## Abstract

Confocal microscopy is an invaluable tool for 3D imaging of biological specimens, however, accessibility is often limited to core facilities due to the high cost of the hardware. We describe an inexpensive do-it-yourself (DIY) spinning disk confocal microscope (SDCM) module based on a commercially fabricated chromium photomask that can be added on to a laser-illuminated epifluorescence microscope. The SDCM achieves strong performance across a wide wavelength range (~400-800 nm) as demonstrated through a series of biological imaging applications that include conventional microscopy (immunofluorescence, small-molecule stains, and fluorescence in situ hybridization) and super-resolution microscopy (single-molecule localization microscopy and expansion microscopy). This low-cost and simple DIY SDCM is well-documented and should help increase accessibility to confocal microscopy for researchers.

## 1. Introduction

Confocal microscopy is a workhorse tool in biological imaging owing to its robust ability to provide high-contrast, 3-dimensional images for a wide range of biological specimens.[1,2] However, commercial confocal microscopes are prohibitively expensive for most individual labs to purchase and maintain, and are thus often limited to core facilities. On the other hand, many labs may already have access to more common epifluorescence or total internal reflection fluorescence (TIRF) microscopes that are equipped with laser sources and sensitive cameras. Here, we describe an alternative solution to bring confocal microscopy within reach for individual research groups by means of a low-cost and easily accessible do-it-yourself (DIY) confocal module that can be straightforwardly added to a laser-illuminated epifluorescence/TIRF microscope.

A popular implementation of confocal microscopy is the spinning disk confocal microscope (SDCM).[3] The spinning disk (SD) at the core of the microscope is a rotating pinhole array placed in the microscope image plane that simultaneously generates an array of excitation point sources while also acting as their conjugate pinholes to reject out of focus light emitted by fluorophores on the sample. Each pinhole within the array sweeps an illumination arc across the sample and the sum of the transmitted fluorescence from these sweeps is integrated by a camera to form an image. Parallelized acquisition then allows the SDCM to achieve high imaging speeds while also causing less photobleaching or phototoxicity than single-point illumination due to lower peak illumination power density for the hundreds to thousands of spots used by the SDCM.[4] The high speed, lower illumination power density, and general convenience of a SDCM comes at the cost of pinhole-pinhole crosstalk that limits optical sectioning performance for thicker samples as compared to point-scanning confocal microscopy where pinhole-pinhole crosstalk does not occur.[5]

A major challenge when implementing SDCM is its inefficient usage of excitation light, the vast majority of which (typically ~95%) is rejected by the SD, a value approximately determined by the fill factor (pinhole area/total area) of the pinhole array. While originally conceived for use with bright reflected light, [6] modern commercial SDCM modules built for demanding fluorescence applications typically use a microlens array that is coupled to the spinning disk in order to focus and concentrate the excitation light onto the SD pinhole array, with a thin dichroic mirror inserted between these spinning elements.[7,8] The microlens array can straightforwardly boost the SD illumination throughput to above 50%, but comes at the expense of higher complexity and cost, a more limited set of pinhole options, and the requirement to use very thin, nonstandard reverse-band (low-pass) dichroic beamsplitters between the microlens and pinhole arrays. In contrast to the early days of SDCM, relatively powerful laser sources (>50 mW) are now commonly available on TIRF and epifluorescence microscopes. These sources are sufficiently powerful to compensate for pinhole losses in order to enable the practical use of low-cost, microlens-free pinhole arrays compatible with the use of standard (high-pass) dichroic beamsplitters.

Here, we describe a microlens-free, do-it-yourself (DIY) approach to building a SDCM using low-cost components on a commercial epifluorescence microscope chassis. While homebuilt SDCMs have been previously reported [9–12] we provide a detailed protocol using modern hardware aimed at researchers familiar with building optical instrumentation. The spinning disk pinhole array at the heart of the DIY SDCM can be affordably fabricated by a photomask manufacturer, and we use a pinhole array design that includes user-determined custom sectors that are optimized for several different applications and that are robust to disk offset (centering). We demonstrate that this microscope is capable of 5-channel imaging spanning the visible and near infrared (~400-800 nm) and we show that it can be used for dim samples that are not typically suited for point-scanning confocal imaging. Finally, with suitable choice of pinhole array parameters and input illumination, we demonstrate super-resolution single-molecule localization microscopy (SMLM) by DNA points accumulation for imaging in nanoscale topography (DNA-PAINT),[13] as well as single-molecule photoswitching-based SMLM by SDCM.[14,15]

## 2. Methods

### 2.1 Design considerations for pinhole array

#### 2.1.1 Details of existing microscope

The epifluorescence/TIRF microscope used as a base in this work was originally constructed for single-molecule localization experiments, and has been described previously.[16] Briefly, the microscope uses five laser lines: a 405 nm line (Coherent, Obis 405nm LX 200 mW) is controlled by direct modulation; the 488 nm (Coherent, Genesis MX488-1000), 561 nm (MPB Communications, 2RU-VFL-P-2000-560-B1R), and 647 nm (MPB Communications, 2RU-VFL-P-1500-647-B1R) lines are modulated by an acousto-optic tunable filter (Crystal Technologies); and a near-infrared 750 nm line (MPB Communications, 2RU-VFL-P-500-750-B1R) is controlled by a filter wheel (Thorlabs, FW102C) with three decades of neutral density (ND) attenuation. *Z*-scanning was performed by an objective nano positioner (Mad City Labs, NanoF100S). Laser modulation and *z*-scanning voltages were provided by two data acquisition cards (National Instruments, PCI-6229 and PCIe-6323) synchronized to the fire signal of the sCMOS camera (Hamamatsu, OrcaV3).

#### 2.1.2 SDCM module

A schematic of the SDCM module and pinhole array are shown in **Fig. 1**. The input beam consisted of the five coaligned excitation sources that were collimated with a ~4 mm diameter and passed through a refractive square shaper (TOPAG, GTH-5-250-4-VIS) to produce a square beam of approximately uniform intensity.[17] For large-area imaging with most general applications, the flattened and square output beam was expanded to ~8 × 8 mm^2^ using a 1:2 telescope (lenses L_1_ and L_2_ in **Fig. 1a**). For small-area imaging with high-intensity illumination, the flattened and square output beam was reduced to ~4 × 4 mm^2^ by omitting lenses L_1_ and L_2_ or further reduced to ~2 × 2 mm^2^ by configuring lenses L_1_ and L_2_ as a reducing 1:2 telescope (see **Fig. 1b-c**). The excitation beam was then relay-imaged to the SD by a pair of lenses (L_3_ and L_4_). The SDCM dichroic mirror was placed in the infinity space of the emission path requiring that one of the relay lenses was shared by both excitation and emission paths. A filter wheel (Thorlabs, FW102C) was used to select the emission filter. For a full parts list and alignment tips, refer to **Table S1** and **Supplemental Note 1. Table S2** includes details on the hardware configuration and illumination conditions for all data sets.

**Figure 1.**
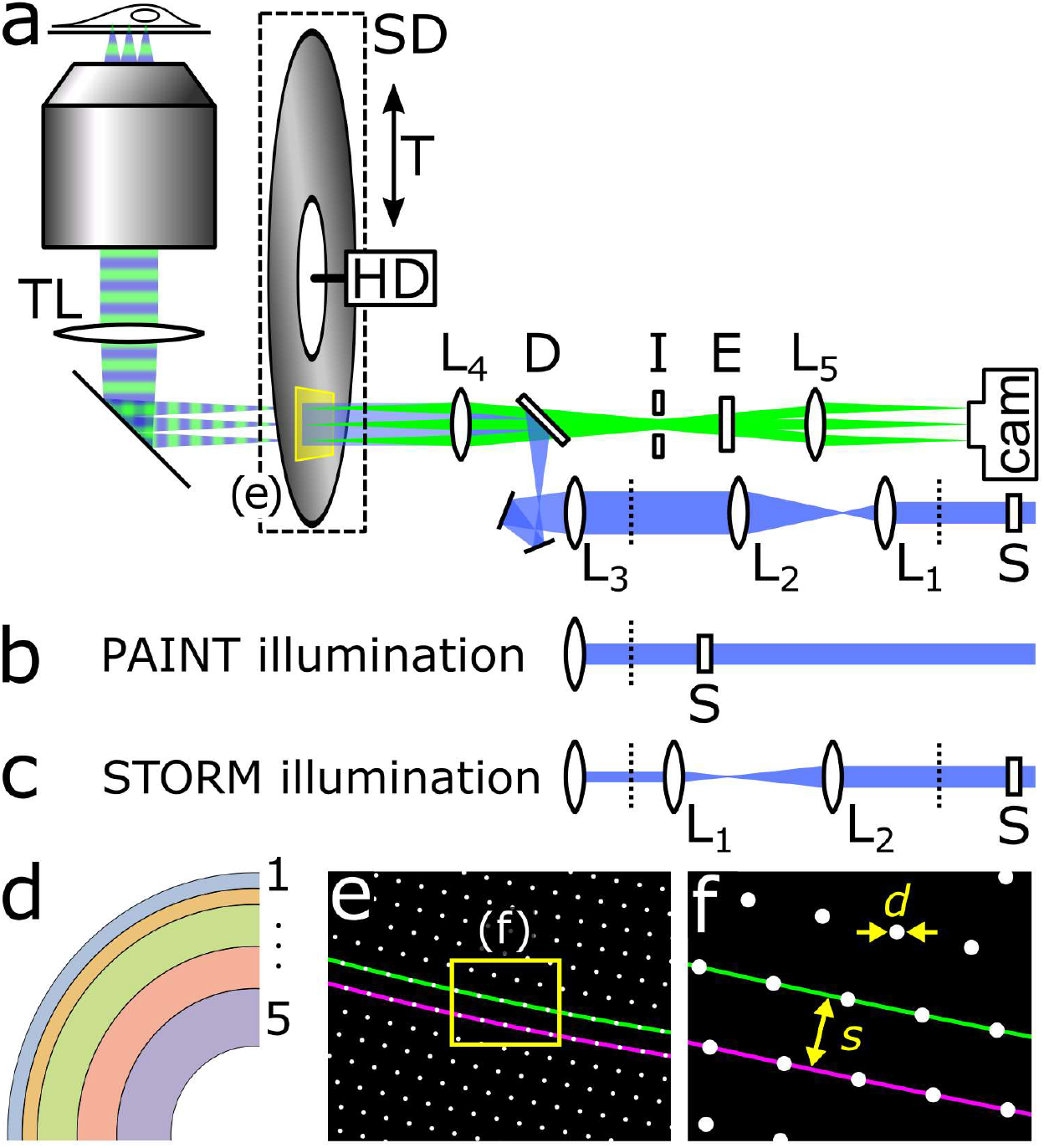
Schematics of spinning disk confocal microscope, disk sectors, and pinhole array. a) Layout of the SDCM with excitation path colored in blue and emission path in green. The blue excitation beam is passed through a refractive square shaper (S) that produces a square, top-hat profile at the adjacent dashed line and is then relay-imaged to the spinning disk (SD) by lens pairs L_1_, L_2_ and L_3_, L_4_. The disk is rotated by a computer hard drive motor (HD) that is mounted on a translation stage (T) for selecting the disk sector or for bypassing the disk (epifluorescence mode). The objective lens (OL) and tube lens (TL) are housed in a commercial microscope chassis. Other listed components are the dichroic mirror (D), iris (I), emission filter (E), and camera. Alternative illumination paths for b) DNA-PAINT by repositioning S and removing L_1_ and L_2_ to achieve no magnification, and for c) SMLM by repositioning lenses L_1_ and L_2_ to achieve two-fold demagnification of the excitation beam, rather than two-fold magnification shown in a). d) Schematic of the five disk sectors used in this work. The narrow sectors 1 and 2 were designed for small area, high power density single-molecule measurements. The remaining sectors were intended for conventional imaging with 100× (3), or 60× immersion objectives (4 and 5). The parameters for each sector are listed in **Table 1**. e) Schematic of a selected area of the sector 3 pinhole array highlighting two neighboring Archimedean spiral paths (green and magenta). f) Zoom in view of the small area in e) with pinhole diameter d and inter-spiral spacing s for the case of s=5×d.

#### 2.1.3 Fabrication of Spinning Disk

Detailed considerations on the design of spinning disk are provided in section 3.1, below (see also **Fig. 1d-f**). The mask was fabricated by Front Range Photomask (Lake Havasu City, AZ) from a 5” × 5” × 0.09” quartz substrate, coated with a ND 5 attenuating chrome layer in all positions except for the pinholes, and cut to 4” outer diameter and with a 0.5” inner diameter hole drilled in the center of the disk. To generate the pinhole array, positions along the concentric Archimedean spirals were calculated using software written in Mathematica and subsequently used to generate a computer-aided design (CAD) file for the photomask vendor. Parameters for the mask direct write process were a 10 µm critical dimension resolution and darkfield pattern polarity. Refer to **Code 1** and **Code 2** for detailed procedure to generate the photomask and the resulting files used in this work.

**Table 1.** Summary of design parameters for multi-sector spinning disk including inner radius *r*_start_, outer radius *r*_stop_, pinhole diameter *d*, inter-spiral spacing *s*, number of concentric spirals *n*, and calculated fill factor (𝜋*d*^2^/4*s*^2^).

| sector | $r_{\text{start}}$<br>[mm] | $r_{\text{stop}}$<br>[mm] | $d$<br>[$\mu\text{m}$] | $s$<br>[$\mu\text{m}$] | $n$ | fill<br>factor | primary use |
| --- | --- | --- | --- | --- | --- | --- | --- |
| 1 | 47.30 | 50.30 | 65 | 163 | 96 | 0.13 | 100× 1.45 NA oil SMLM |
| 2 | 44.25 | 47.25 | 65 | 244 | 32 | 0.056 | 100× 1.45 NA oil SMLM |
| 3 | 36.20 | 44.20 | 50 | 250 | 32 | 0.031 | 100× 1.45 NA oil |
| 4 | 28.15 | 36.15 | 35 | 350 | 12 | 0.0079 | 60× 1.27 NA water lens<br>("double wide") |
| 5 | 17.10 | 28.10 | 35 | 175 | 32 | 0.031 | 60× 1.27 NA water lens<br>20× 0.45 NA air lens |

For the spinning disk motor, a stripped-down hard drive (Seagate ST500DM002) motor-base assembly was modified by machining a window to provide a light path. The pinhole array was secured onto the spindle motor by a machined Delrin adapter (**Fig. S1** and **Code 3** for CAD files). Note that the adapter was designed for disks with 0.09” glass thickness and uses two inset O-rings to gently, but securely grip the glass disk. It was important for the disk to be oriented so that the reflective (metallic) side of the disk was facing the light source. The motor-base assembly was mounted to an aluminum base and translation stage (Newport, PRL-6) that allowed manually selecting different sectors of the disk, or complete bypass of the disk for epifluorescence imaging. The hard drive speed was controlled by a three-phase brushless direct-current (DC) motor controller (Yeeco, ZS-X9B) powered by a 5 V power supply. A microcontroller (Arduino, Uno) and a reflective optical switch measured the real-time readout of the disk rotation rate (**Fig. S2** and **Code 4**). The disk was run consistently at ~4560 RPM (rotations per minute). For a detailed list of all hardware components refer to **Table S1**.

### 2.2 Sample preparation

#### 2.2.1 Chemicals and reagents

Primary antibodies were purchased as follows: rat anti-alpha tubulin (MA1-80017, Thermo Fisher Scientific), mouse anti-alpha tubulin (62204, Thermo Fisher Scientific), rabbit anti-detyrosinated tubulin (ab48389, Abcam), mouse anti-vimentin (18-0052, Invitrogen), rabbit anti-homer1 (Synaptic Systems, 160003) and mouse anti-bassoon (Abcam, ab82958). Unconjugated donkey anti-rat (712-005-153), donkey anti-rabbit (711-005-152), and donkey anti-mouse (715-005-151) secondary antibodies were purchased from Jackson ImmunoResearch. Bovine serum albumin (BSA; BAS-50) was purchased from Rockland Immunochemicals Inc. Fluorescent dyes and biotin were purchased as follows: Alexa Fluor 750 NHS ester (A20011), Alexa Fluor 647 NHS (A20006), Alexa Fluor 568 NHS ester (A2002), Alexa Fluor 488 NHS ester (A20000), Alexa Fluor 405 NHS ester (A30000) and EZ-Link NHS-PEG4-Biotin were purchased from Thermo Fisher. ATTO 647N NHS ester (18373-1MG-F), ATTO 565 NHS ester (72464-1MG-F) and ATTO 488 NHS ester (41698-1MG-F) were purchased from Sigma Aldrich. Other reagents included: Glutaraldehyde (50%, GA, 16320) and paraformaldehyde (32%, PFA, RT15714) from Electron Microscopy Sciences; Acrylamide (40%, 1610140) and bis-acrylamide (2%, 14101420) from Bio-Rad Laboratories; Ammonium persulfate (APS, 17874), tetramethylethylenediamine (TEMED, 17919) and proteinase K (EO0491) from Thermo Fisher Scientific; Sodium acrylate (408220), Hoechst 33258 (B2883-25MG), 4-hydroxy-2,2,6,6-tetramethylpiperadin-1-oxyl (TEMPO, 176141), guanidine hydrochloride (G3272), methacrylic acid NHS (MA-NHS, 730300), piperazine-1,4-bis(2-ethanesulfonic acid) disodium salt (PIPES disodium salt, P3768), poly-l-lysine (P8920), 6-Hydroxy-2,5,7,8-tetramethylchromane-2-carboxylic acid (Trolox, 238813), glucose oxidase (G2133-50KU), and catalase (C100) from Sigma-Aldrich.

#### 2.2.2 Resolution measurement specimens

Point-spread function (PSF) measurements for 488 nm, 561 nm and 647 nm excitation were conducted using 0.1 µm TetraSpeck beads (Thermo Fisher, T7279) immobilized on coverglass. The coverslip was rendered hydrophilic by exposure to an air plasma for 1 min, then 10 µL of a 1:50 dilution of beads in ethanol were dried on the surface. For the high numerical aperture (NA) air objective (Nikon, 100× 0.9 NA, MUE13900), beads were imaged directly on the coverglass. For the water-immersion objective (Nikon, 60× 1.27 NA, MRD07650) the immobilized beads were submerged in water, while for the oil-immersion objective (Nikon, 100× 1.45 NA, MRD01905) the beads were embedded in optical cement (Nordland, NOA 60). For the embedded beads, a bead-coated coverglass was inverted onto 5 µL of optical cement and cured with a 15 s exposure from a UV transilluminator. Immersion oil with *n* = 1.512 was selected from a set ranging from 1.500 – 1.530 (Cargille, Laser Liquids) for imaging. For the low NA air objective (Nikon, 20× 0.45 NA, MRH38220), 0.2 µm TetraSpeck beads were used.

Concentrated dye measurements were performed using a ~1 M aqueous fluorescein solution (Fisher, S25328). A thin channel was formed between a #1.5 coverglass and a microscope slide by two strips of double-sided tape, then filled with dye solution, and sealed with epoxy.

#### 2.2.3 Sample preparation for cultured cells

Cell culture samples for PSF measurements, 5-channel imaging, DNA-PAINT, or STORM experiments were prepared using minor modifications to a standard immunofluorescence protocol.[18] Briefly, approximately 40,000 cells (BSC-1, PTK-1 or hTERT RPE-1) were seeded per well in an 8-well coverglass chamber (Ibidi, 80827) and allowed to adhere overnight. The cells were extracted for 30 s (0.5% Triton X-100 (TX-100), 100 mM PIPES), fixed for 10 min (3.2% paraformaldehyde (PFA), 0.1% glutaraldehyde (GA), 100 mM PIPES), and rinsed with excess phosphate-buffered saline (PBS). Blocking buffer (3% bovine serum albumin, 0.5% TX-100, PBS), was used for all subsequent staining steps involving antibodies, streptavidin, or DNA, followed by washing with PBS for 30 min. The sample was blocked for 1 hour, followed by incubation with primary antibodies (1-2 µg/mL) overnight, labeled with secondary antibodies (5 µg/mL) for one hour, and then postfixed with 0.1% GA in PBS for 10 min. Small-molecule stains such as Hoechst 33342 (1 µg/mL) and Alexa Fluor 647 Phalloidin (Life Technologies, 0.5 μM) were applied immediately prior to imaging for 10 min in PBS. For DNA-PAINT samples, a biotin modified secondary antibody (2 µg/mL) and a dilute ATTO 488 labeled secondary antibody (0.2 µg/mL) were used, followed by application of unlabeled streptavidin (5 µg/mL) for 30 min, and finally a 500 nM solution of biotin-modified P1 docking sequence for 30 min in blocking buffer containing 0.1% TX-100 and 1 mg/mL yeast tRNA. The small quantity of ATTO 488 labeled antibody aided in focusing for DNA-PAINT imaging. DNA-PAINT reporter sequence P1 labeled with Cy3B (sequence P1) was a gift from B. Beliveau. Immunofluorescent samples were imaged in an oxygen-scavenging buffer (Glox, 100 mM Tris pH 8, 10% glucose (wt/wt), 0.5 mg/mL glucose oxidase, 40 µg/mL catalase) with 1 mM trolox. For DNA-PAINT samples, Glox was supplemented with 250 mM NaCl, 0.1% Tween 20, and 0.05 nM of sequence P1-Cy3B. For STORM samples, Glox containing 143 mM 2-mercaptoethanol was used for photoswitching, while for 3D STORM the buffer also included 60% sucrose to boost the refractive index and limit spherical aberration.[19] Refer to **Table S2** for detailed staining and imaging conditions and **Table S3** for DNA sequences.

#### 2.2.4 Sample preparation for mRNA FISH labeling of expanded cultured cells

Single-molecule mRNA FISH imaging was conducted on samples prepared with a slightly modified ExM and FISH protocol. Approximately 70,000 RPE-1 cells were seeded on a 12 mm diameter round coverglass within a well of a 24 well plate and allowed to adhere overnight. The cells were briefly washed in PBS and then fixed in 4% PFA in PBS for 10 min. The cells were extracted in 70% ethanol for 10 min and then equilibrated in wash buffer (2× saline-sodium citrate (SSC), 10% formamide) prior to use. The sample was treated with a 50 µL of a 125 nM probe set specific to mRNA for *GAPDH* (glyceraldehyde 3-phosphate dehydrogenase) in wash buffer containing 10% dextran sulfate overnight at 37 °C and then washed three times for 10 min in excess wash buffer. The sample was treated with a 25 mM methacrylic acid methacrylic acid N-hydroxysuccinimidyl (MA-NHS) ester solution in PBS, and then gelled as described below. Gelation was performed by incubating the coverslip containing the sample in monomer solution (1 × PBS, 2 M NaCl, 2.5% acrylamide 0.15% N,N-methylenebisacrylamide, 8.625% sodium acrylate) for 5 min. Meanwhile, 70 µL of monomer containing 0.2% ammonium persulfate and 0.2% tetramethylethylenediamine was placed onto a hydrophobic glass surface and the coverslip was inverted onto the droplet and polymerized in a N_2_ atmosphere for 15 min. The sample was allowed to digest overnight in digestion buffer (1× Tris-acetate-EDTA, 0.8 M guanidine hydrochloride, 0.1% TX-100) containing 1% proteinase K. The sample was washed in a large excess of 2× SSC, and then 10 nM of ATTO 565 labeled oligonucleotide readout probe was allowed to hybridize overnight. The sample was washed in a large excess of 2× SSC to remove unhybridized readout probe, 1 µM of TO-PRO-3 iodide (Thermo Fisher, T3605) was added for 30 min, and then the sample was expanded in 0.01× SSC until it reached ~3× expansion. The sample was mounted on a poly-l-lysine coated coverglass for imaging.

The GAPDH probe set contained 24 targeting sequences, each of 26-32 nts in length, and was ordered from IDT each normalized to 10 mM concentration.[20] Using an equal component mixture of all sequences, the probe set was enzymatically modified with an amino-11-ddUTP (Lumiprobe, A5040) using a terminal transferase (New England BioLabs, M0315S) as described previously.[21] Briefly, 30 µL of the mixed probe set probe set (10 mM total DNA) was reacted with a 3-fold molar excess of ddUTP using 20 units of enzyme following the enzyme manufacturer’s instructions. The modified probe set was then used without further purification and stored at -20 °C. The ATTO 565 labeled reporter oligonucleotide was purchased from IDT. Refer to **Table S3** for GAPDH probe set sequences.

#### 2.2.5 Expanded brain tissue

Expanded brain tissue was processed as detailed previously.[22,23] Briefly, mice (genotype Glt25d2-Cre_NF107;Ai14) were anesthetized with isoflurane and perfused transcardially with PBS, followed by paraformaldehyde (4% PFA in PBS). The brain was removed by dissection, postfixed in 4% PFA in PBS for 3-6 hr at room temperature and then overnight at 4 °C. Fixed brains were washed in PBS and stored in PBS with 0.02% sodium azide prior to sectioning 100 µm thick coronal slices using a vibratome. Slices were soaked in blocking buffer for 6 hours, incubated with primary antibodies in blocking buffer for 24 hr, and then incubated with secondary antibody in blocking buffer for an additional 24 hr. The sample was kept at 4 °C for the duration of the staining and was washed three times in PBS for 20 min after each antibody step. The brain slices were treated with 1 mM MA-NHS in PBS for 1 hr at room temperature and then equilibrated in monomer solution for 1 hr. The tissue was then gelled while sandwiched between two coverglass using the same recipe as above, but with the addition of 0.01% of the inhibitor 4-hydroxy-2,2,6,6-tetramethylpiperidin-1-oxyl (4-hydroxy-TEMPO) under a humidified N_2_ atmosphere at 37 °C for 3 hr. The sample was digested overnight as described above, expanded in water, and mounted on a poly-l-lysine coated coverglass for imaging. Refer to **Table S2** for detailed staining and imaging conditions.

### 2.3 Data acquisition and analysis

All SDCM instrumentation and data acquisition was controlled by the open-source software Micromanager.[24] The acquisition order was *z*-scan, then wavelength in order to take advantage of the hardware timed *z*-scanning piezo and minimize the overhead of the filter wheel movement. All image analysis was conducted using routines written in Mathematica and Python. Bead PSFs were fit to 3-dimensional Gaussian profiles, whereas lateral and axial profiles of microtubules were fit to 2-dimensional Gaussian profiles. Expanded brain data was tiled in Micromanager, stitched using BigStitcher,[25] and rendered with 3Dscript.[26] Two-dimensional localization microscopy data (STORM and DNA-PAINT) were analyzed as described previously.[27] Three-dimensional STORM data was analyzed and rendered using ThunderSTORM,[28] and the resulting images were registered by rigid alignment using Elastix.[29]

## 3. Results

### 3.1 Design considerations and theory

A SDCM uses an array of pinholes on a disk located at the conjugate image plane to create an array of excitation beams and detection pinholes that scan the sample when the disk is rotated (**Fig. 1a**). This parallelization can increase the imaging speed up to 2-3 orders of magnitude as compared to point-scanning confocal microscopy with a single excitation spot (~1 Hz).[8] However, simultaneous excitation points also permit pinhole-pinhole crosstalk, where out of focus light passes through neighboring pinholes and limits optical sectioning performance for thicker specimens.[30] The specific design of the disk pinholes involves making tradeoffs between optical sectioning, scan speed, efficiency of transmission of excitation light, and efficiency of transmission of emission light.

Prior work has established some general guidelines for design of pinhole arrays in a SD [30–35]. First, the pinhole diameter *d* is typically chosen to be approximately equal to the diameter *w* of the first minimum of an Airy disk PSF given by [1]

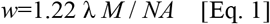

where λ is the wavelength of light, *M* is the magnification between the sample and SD, and *NA* is the numerical aperture of the objective lens. Setting the pinhole diameter *d* to equal the Airy diameter *w*, which corresponds to 1 Airy unit, provides a good balance of rejecting out-of-focus light and efficient transmission of in-focus light.[33] For λ=600 nm, a 60× 1.27 NA water-immersion objective lens yields *w*=34 µm and a 100× 1.45 NA oil-immersion objective lens yields *w*=50 µm.

A second major consideration is the placement of pinholes on the SD. Pinholes are typically placed along one or more Archimedean spirals, spaced by a constant arclength that is also equal to the inter-spiral spacing (**Fig. 1e-f)**.[8] In this arrangement, at a distance sufficiently far from the center, the pinholes form a grid-like pattern (**Fig. 1e**). Additionally, this distribution of pinholes maintains an approximately uniform hole density at different distances from the disk center to ensure even transmission at all regions of the disk. Third, the inter-spiral spacing *s* is typically selected to be ~3-10 times the pinhole diameter *d* (**Fig. 1f**), such that the resulting fill factor, or fraction of the disk that contains holes, is approximately given by 𝜋*d*^2^/4*s*^2^ and typically has a value of ~1-10%.[1] This leads to a reasonable tradeoff between fill factor, which approximately equals the efficiency of transmission of the illumination light in our microlens-free disk, and pinhole separation, which enables low pinhole-pinhole crosstalk for sufficiently thin specimens.

While a single spiral of pinholes is in principle suitable for a SD, the use of multiple concentric spirals leads to greatly decreased sensitivity of the illumination uniformity to offset of the disk from its true center position (**Fig. 2**). This issue of sensitivity to offset appears to have received little attention in the scientific literature, although multi-spiral designs are commonly used.[3,36] A set of *n* concentric Archimedean spirals in polar coordinates is given by Equation 2

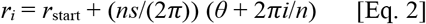

where *r*_*i*_ is the radius of the *i*^th^ spiral (indexed from *i*=0 to *n*-1) as a function of angle *θ, s* is the inter-spiral spacing, and *r*_start_ is the radial position of the innermost pinhole. Although all pinhole patterns tested in the laboratory used patterns with pinhole arclength spacing equal to inter-spiral spacing *s*, it is convenient to simulate the sensitivity to offset by considering the case of a very small pinhole arclength spacing (**Fig. 2b**). There we consider *n*=1, 2, 4, 8, 16, or 32 spirals, *r*_start_=40,000 µm, inter-spiral spacing *s*=250 µm, and pinhole arclength spacing 2.5 µm. A single spiral would perform poorly (normalized root mean square deviation (RMSD) =0.14 for illumination intensity) with even just 10 µm offset, which is an extremely difficult tolerance to achieve experimentally, while a 32-spiral design would perform well with an offset of up to ~1000 µm (normalized RMSD=~0.003), a tolerance that is very easily achieved in practice. Similar results were obtained in a more detailed simulation based on disk patterns we used in the laboratory where inter-spiral spacing *s*=250 µm is equal to the pinhole arclength spacing (**Fig. 2c-d**).

**Figure 2.**
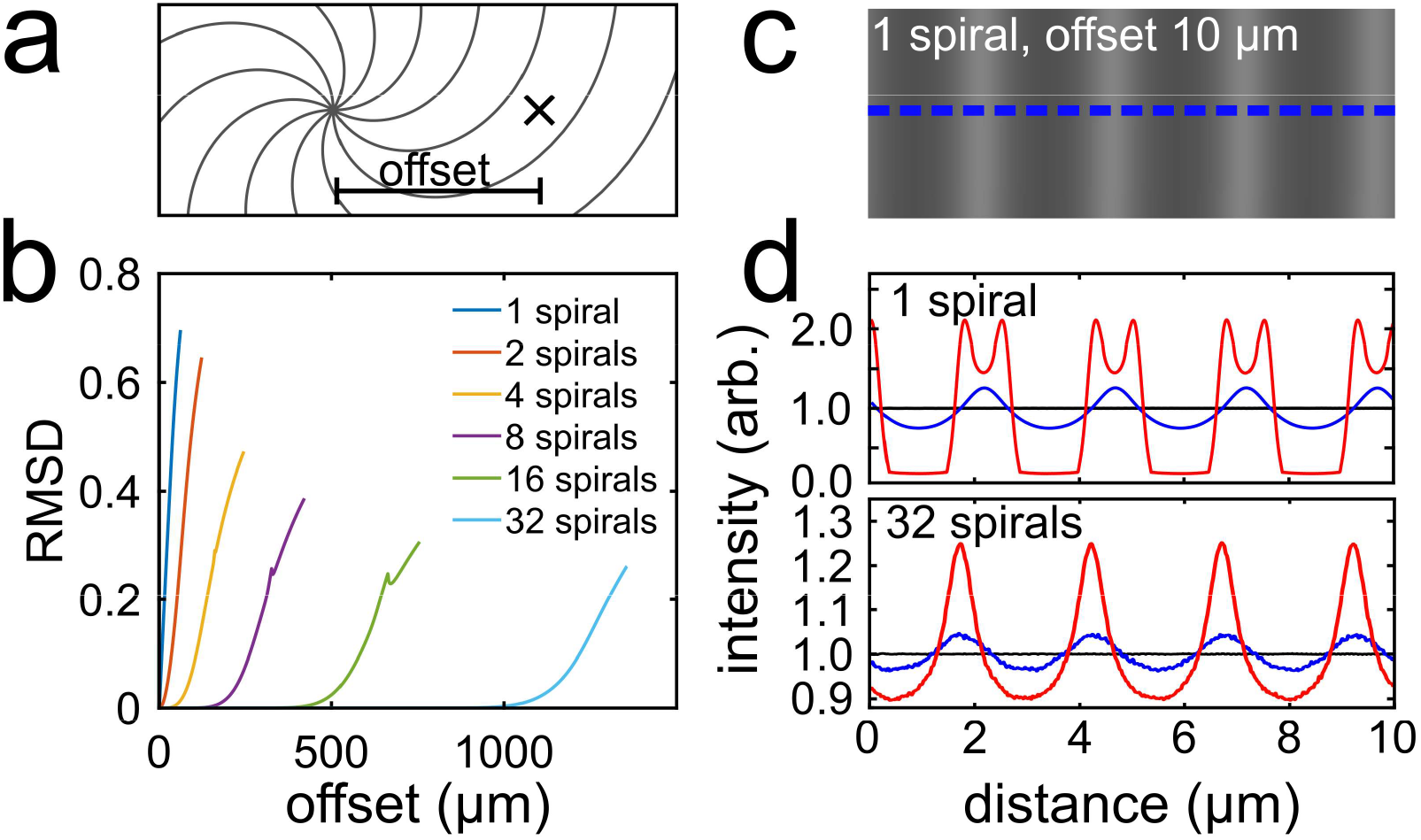
Simulated spinning disk illumination intensity patterns for misalignment offset. a) Schematic diagram of the misalignment offset between the Archimedean spiral origin and the actual rotational axis of the disk denoted by the cross. b) Simulated root mean square deviation (RMSD) of illumination intensity vs disk offset from the center for patterns consisting of 1-32 concentric spirals. All patterns used a constant inter-spiral spacing of 250 µm, but a pinhole arclength spacing of 2.5 µm at a radius of ~40 mm from the spiral center, where 1 spiral completed 32 revolutions, 2 spirals completed 16 revolutions, etc. c) Simulated illumination intensity for a single spiral disk with an offset of 10 µm in a 10 µm × 4 µm area of the image plane. d) Simulated illumination intensity profiles for the single spiral disk in (c) with offsets of 0 µm (top, black), 10 µm (top, blue) and 100 µm (top, red), and for a 32-spiral disk with offset of 0 µm (bottom, black), 1100 µm (bottom, blue) and 1200 µm (bottom, red). The intensity profiles were normalized to the 0 µm offset intensity level. The simulations in (c) and (d) used pinhole diameter d = 50 µm and inter-spiral spacing s=250 µm, at a radius of ~40 mm from the spiral center, and were simulated as viewed by a 100× lens.

In principle, a very large number of concentric spirals could be used while still maintaining the inter-spiral distance to be equal to *s*, but the radial increment of successive pinholes within a spiral should not exceed half the full width at half maximum (FWHM) of the PSF in order to fulfill the Nyquist sampling criterion along the lateral dimension. The radial increment of successive pinholes ∆*r* at an approximate distance of *r*_avg_ from the center for pinholes along *n* concentric Archimedean spirals is given by

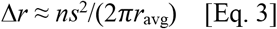

For example, a disk with *n*=32 concentric Archimedean spirals, an inter-spiral spacing *s*=250 µm, and *r*_avg_=40,000 µm from the disk center, has a lateral increment of ∆*r*≈8 µm which is a fraction of the typical Airy disk width (assuming a 100× 1.45 NA objective), indicating that the lateral sampling is equal to or better than the Nyquist sampling limit required for diffraction-limited imaging.

A third consideration is the inefficient use of excitation light. As described above, modern commercial SDCMs often use a microlens array prior to the pinhole array. A microlens array has the advantage of boosting excitation transmission to >50%, which is more than 16 times higher than is achieved by a microlens-free system with a typical fill factor of, for instance, 0.031, but comes at the cost of higher complexity, higher cost, and the necessity of using small and thin reverse-band (e.g., short-pass) dichroic filters positioned between the microlens array and SD. Fortunately, high-powered continuous-wave lasers have become sufficiently cheap that microlens-free systems are feasible and can largely compensate for these losses. In addition, to best utilize the available laser power, we incorporated into the illumination path a refractive beam shaper [17] to convert the approximately Gaussian laser profile into a square beam of nearly uniform intensity as well as an optional 1:2 magnifying telescope or a 2:1 demagnifying telescope (**Fig. 1a-c**). These measures enabled sufficient illumination intensity with use of a microlens-free SD design for either low to moderate intensity illumination of large-areas (for conventional imaging of cell and tissue specimens) or for relatively high-intensity illumination of smaller areas (for SMLM).

Based on the above several considerations for pinhole diameter, inter-spiral spacing, the use of multiple concentric Archimedean spirals, and the microlens-free approach, we designed five multi-spiral pinhole arrays that were compatible with the objective lenses available on our microscope and which are suitable for different experiments. These pinhole arrays, summarized in **Table 1**, were designed to occupy 5 different radial sectors of a disk (**Fig. 1d**) so that the disk can be translated within the plane of the disk to enable use of different pinhole arrays without the need to exchange the disk. Sectors 1-2 (**Table 1**) were designed for high intensity imaging. The remaining three sectors were designed for conventional imaging for a 100× 1.45 NA oil-immersion objective (sector 3) and a 60× 1.27 NA water-immersion objective (Sectors 4-5). As described in the methods, the SD pinhole array was fabricated to order from a glass or quartz photomask (typically used for photolithography) that had been cut and drilled to a disk shape. Photomasks offer a customizable, accessible, low-cost, and high-resolution solution, thus making the DIY SDCM feasible. Basic or prototype disks can be purchased for ~$400, while the design used for most measurements in this work incorporated an additional broadband anti-reflection coating raising the total cost of our five-sector disk to ~$1000.

### 3.2 Incorporation into existing microscope

We incorporated the SDCM module into an existing epifluorescence/TIRF setup based on a Nikon Ti-U microscope chassis. The microscope is configured with epifluorescence/TIRF illumination from the rear port and relies on a dichroic beamsplitter in the microscope filter cube residing in the infinite path between the objective and tube lens with the left microscope port used for detection. The SDCM module was incorporated into the right-side port of the microscope and intended to be used when the epifluorescence/TIRF dichroic beamsplitter was removed (refer to **Supplemental Note 1** for alignment tips). On top of the existing epifluorescence/TIRF setup, the total cost of additional components for the SDCM module was approximately $7,000, where the majority this cost was accounted for by a pentaband dichroic, an emission filter set, and a motorized filter wheel (**Fig. 1**; see **Table S1** for a detailed list). Custom components included the SD pinhole array and a machined motor assembly adapter.

To spin the disk, we modified a discarded Seagate hard drive disk motor since the hard drive axle has low axial play, can achieve high speeds (4,000-10,000 RPM), and since the speed can be finely controlled using a brushless DC motor controller. We mounted the SD onto the motor by machining a custom polymer (Delrin) adapter and clamping plate to secure the SD to the hard drive case (**Code 3**). While we have only tested the adapter with a particular model of Seagate spindle motors, it should be possible with minor modifications of the polymer adapter to attach the SD to the spindle motors of other brands. We also designed an aluminum base to mount the motor-disk assembly to an optical table or to a sliding rail for easy selection of the spinning disk sector (**Code 3**).

The disk was carefully aligned onto the motor spindle to be close to the motor axis. We did this by securing the disk to the motor spindle with just enough pressure to prevent slipping but while enabling it to be displaced by a gentle tap from a finger or a light, soft-tipped mallet; in this state, when slowly rotating the unpowered disk by hand, one could observe subtle wobble of the disk and carefully correct the offset by tapping the disk in a suitable direction before tightening the spindle. While the use of multi-spiral disk designs described in **Fig. 2** made the illumination uniformity not particularly sensitive to offset of the disk center, good centering ensured rotational balance. We did not observe noticeable vibrations for a carefully centered disk with our typical rotation speed of ~4560 RPM. However, as a precaution, we created a metal housing around the motor-disk assembly to contain debris in the event of a mechanical failure of the rapidly spinning components and to also protect the disk’s surface from unintentional damage by the user.

### 3.3 Characterization

#### 3.3.1 SDCM Spatial Resolution

We used ~100 nm diameter fluorescent microbeads to measure the PSF for the 100× 1.45 NA objective with disk sector 3 (*d*=50 µm and *s*=250 µm) of the SDCM (**Fig. 3**). With 488 nm excitation, the PSF had a lateral FWHM of ~210 nm and an axial FWHM of ~420 nm; these values were sensitive to immersion oil index of refraction (**Fig. S2**). The lateral and axial PSF values are summarized in **Table 2** for all five excitation channels from 405 nm to 750 nm for sector 3 of the disk (see also **Table S4** for a full list of objective-pinhole combinations evaluated). Note that while PSFs at visible wavelengths (488 nm, 561 nm, and 647 nm) could be robustly measured using sub-wavelength beads, we were unable to obtain high performance beads for 405 nm or 750 nm excitation, and we opted instead to use the transverse and axial profile of immunostained microtubules in cultured cells as an alternative metric for quantifying resolution.

**Figure 3.**
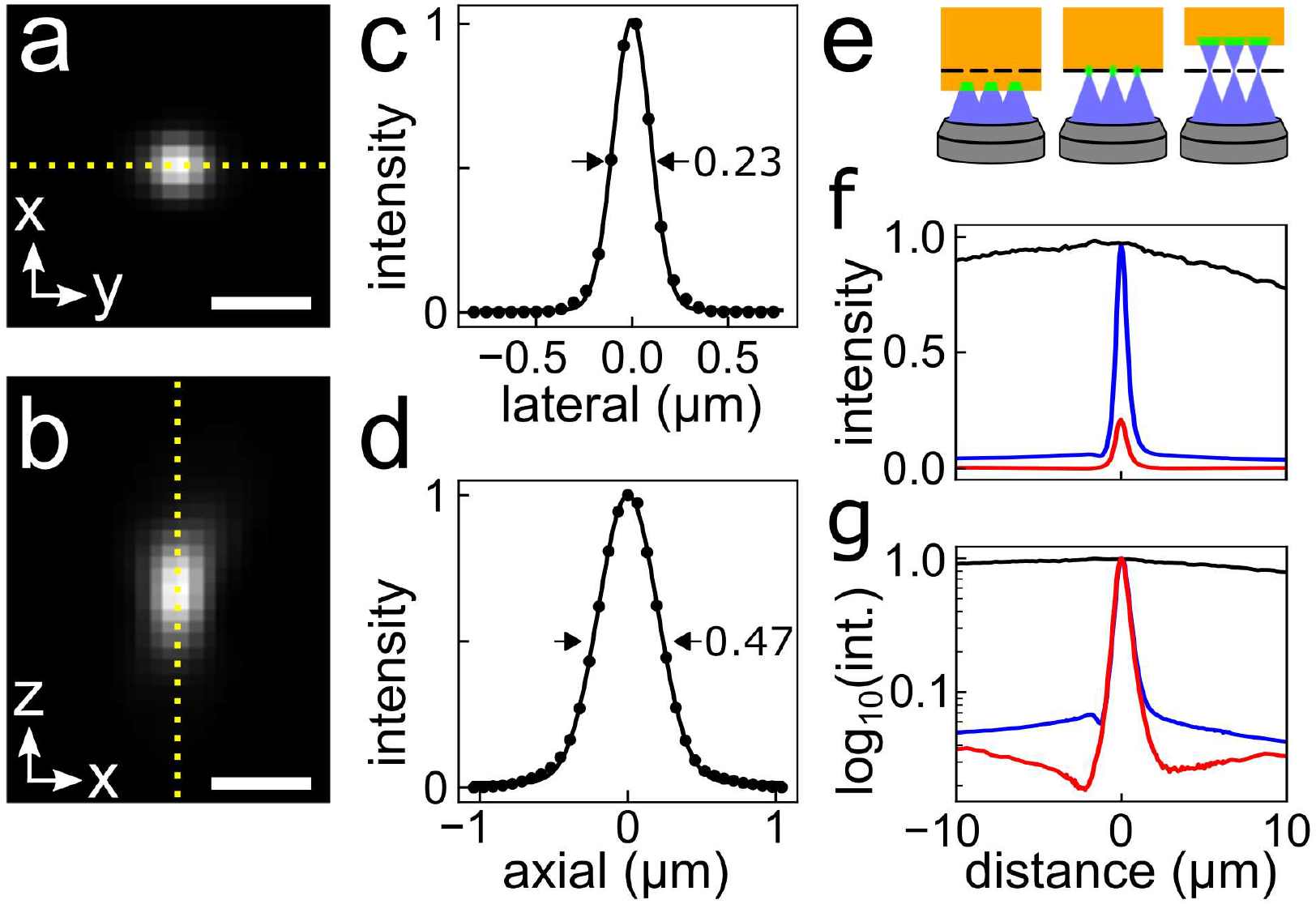
Point spread function (PSF) and optical sectioning with SDCM. a) x-y view and b) x-z view of PSF measurement performed with SDCM using 100 nm fluorescent beads excited at 488 nm. Imaging was performed using a 100× 1.45 NA oil-immersion objective lens and disk sector 3 (**Table 1**). c-d) Lateral and axial profiles of PSF centered on the dashed yellow lines in panels **a** and **b**, respectively, with points showing measured data and lines showing the 3D Gaussian fits. e) Schematic diagram of concentrated dye assay to produce a thin fluorescent plane. The dotted black line represents the virtual image of the pinhole in the sample plane. f) Concentrated dye signal as a function of axial position using a 100× 1.45 NA water-immersion objective lens for epifluorescence (black), SDCM with d = 50 µm and s=250 µm (blue), or SDCM with d=50 µm and s=500µm (red). g) Normalized axial profiles of the concentrated dye signal shown from panel **f** on a Log_10_ plot. Scale bars, 500 nm (a,b).

**Table 2.** Resolution of SDCM with 50 µm pinholes and high NA objective lenses.

| | 100× 1.45 NA Oil | | 60× 1.27 NA H<sub>2</sub>O | | 100× 0.9 NA Air | |
| --- | --- | --- | --- | --- | --- | --- |
| $\lambda_{\text{ex}}$<br>(nm) | $X_{\text{FWHM}}$<br>(nm) | $Z_{\text{FWHM}}$<br>(nm) | $X_{\text{FWHM}}$<br>(nm) | $Z_{\text{FWHM}}$<br>(nm) | $X_{\text{FWHM}}$<br>(nm) | $Z_{\text{FWHM}}$<br>(nm) |
| 405<sup>a</sup> | 219 | 964 | 303 | 1370 | - | - |
| 488<sup>b</sup> | 222 | 487 | 308 | 820 | 317 | 737 |
| 561<sup>b</sup> | 237 | 500 | 329 | 825 | 334 | 809 |
| 647<sup>b</sup> | 273 | 561 | 353 | 845 | 386 | 930 |
| 750<sup>a</sup> | 349 | 604 | 444 | 1044 | - | - |

<sup>a</sup> FWHM (full width at half maximum) values obtained by 2-dimensional Gaussian fit to transverse lateral and axial profiles of immunostained microtubules.
<sup>b</sup> FWHM values obtained by 3-dimension Gaussian fit of multispectral 100 nm fluorescent beads.

#### 3.3.2 SDCM Rejection of out of focus light

We used a concentrated (~1 M) aqueous solution of fluorescein excited with 488 nm illumination to measure the rejection of out of focus light using a prototype disk containing sectors with a ~3% fill factor (*d*=50 µm and *s*=250 µm) and one with ~1% fill factor (*d*=50 µm and *s*=500 µm). The solution is sufficiently concentrated that only a thin layer of dye is excited near the substrate (**Fig. 3e**), but, unlike fluorescent beads, the signal from a thin uniform sheet of dye does not diminish due to spreading out laterally as a function of defocus.[37,38] As expected, the larger fill factor exhibited a brighter signal but less rejection of out of focus light (**Fig. 3f-g**) because its higher density of pinholes more efficiently transmitted excitation light and more efficiently permitted pinhole-pinhole crosstalk.[39] In contrast, epifluorescence images showed approximately constant signal over a ±10 µm scan range (**Fig. 3f-g**).

### 3.4 Applications

#### 3.4.1 SDCM Performance with Conventional and Hydrogel-Expanded Specimens

We tested the performance of the SDCM using a range of specimens. **Fig. 4a-f** shows a composite five-channel image of a single plane of a BS-C-1 cell stained for tyrosinated tubulin, detyrosinated tubulin, actin, vimentin, and DNA that was recorded using a 100× 1.45 NA oil-immersion objective lens with disk sector 3. The system showed good performance across all channels with these relatively bright stains.

**Figure 4.**
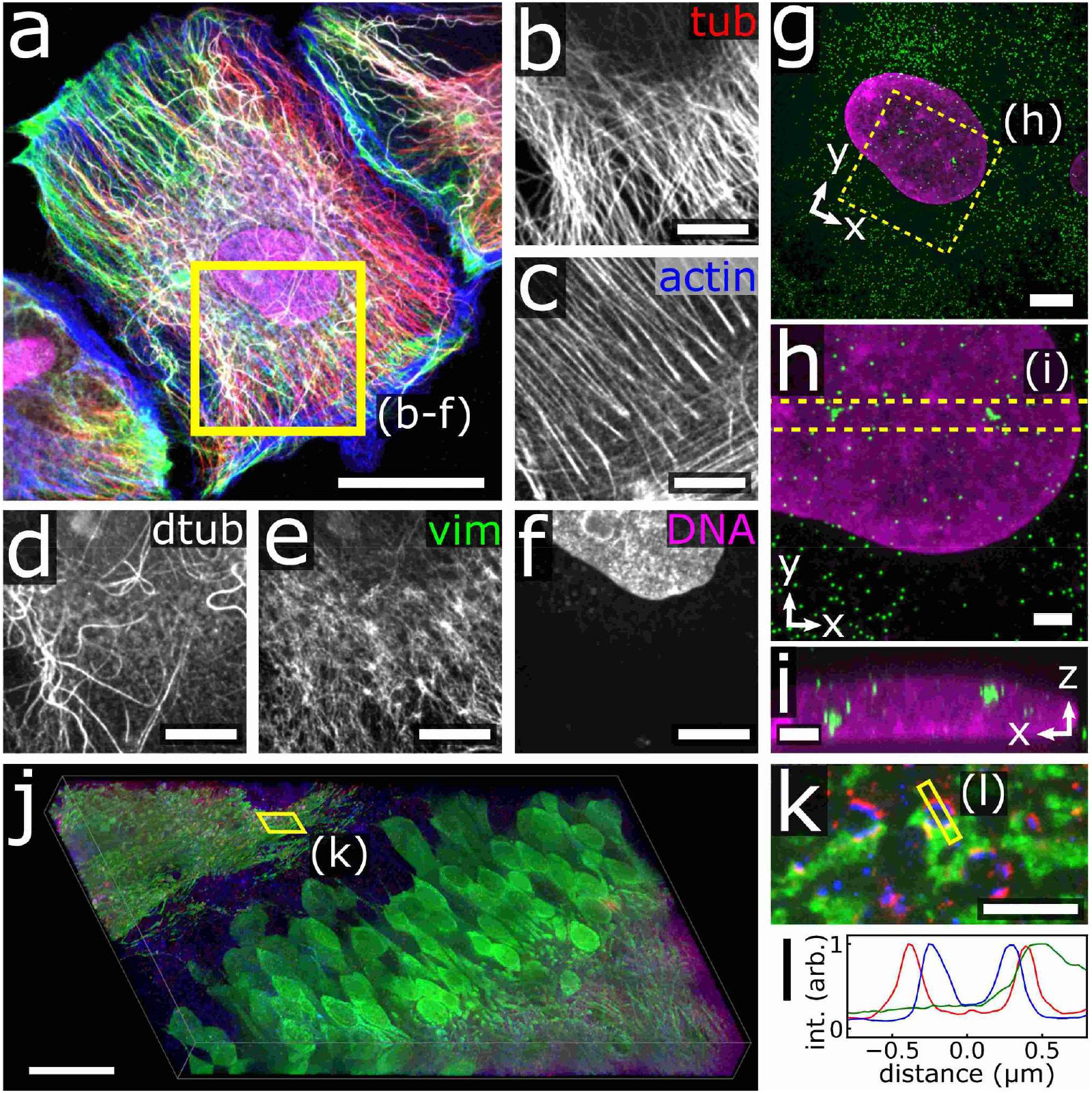
Selected biological specimens imaged with SDC microscope. (a) BSC-1 cells stained for the cytoskeleton and DNA. The individual channels and excitation wavelength of the boxed region in **a** are shown in panels: (b) tyrosinated tubulin (tub, 750 nm), (c) actin (647 nm), (d) detyrosinated tubulin (dtub, 561 nm), (e) vimentin (vim, 488 nm), and (f) DNA (405 nm). (g) Maximum intensity projection of expanded RPE-1 cells stained for GAPDH mRNA (green) and DNA (magenta). (h) Zoom-in of the area highlighted in **g**, showing nascent mRNA clusters, and corresponding transverse projection (i) of the area highlighted in **h**. (j) Expanded brain immunostained for tdTomato (green), Homer (red), and Bassoon (blue) tiled and stitched using SDCM. (k) Zoomed-in view of a single plane of the boxed region in **j**, and corresponding line profile of a synapse in the boxed region in **k**. Scale bars, 30 µm (a), 10 µm (b-f), 5 µm (g), 2 µm (h,i), 25 µm (j) and 2.5 µm (k). Scale bars in expanded samples are given in pre-expansion dimensions.

Next, we prepared a more challenging specimen for SDCM in which individual *GAPDH* mRNA molecules were labeled by fluorescence *in situ* hybridization (FISH) in RPE-1 cells prepared using expansion microscopy (ExM) and expanded isotropically three-fold along each of three axes. In ExM, a swellable hydrogel is synthesized within a fixed specimen and then enlarged upon incubation in deionized water so that features closer than the ~250 nm diffraction limit of light can be resolved in the expanded state.[22,23] Using a 60× 1.27 NA water-immersion objective lens with disk Sector 5 we found that *GAPDH* mRNA were relatively abundant, but were well-resolved thanks to substantial de-crowding achieved by the ~27-fold (volumetric) expansion (**Fig. 4g-i**). Furthermore, we were able to observe the 3-dimensional structure of bright mRNA clusters within the nucleus that presumably label nascent mRNA near *GAPDH* loci (**Fig. 4h-i**).

Finally, we imaged a portion of an expanded mouse brain hippocampus immunostained for a subset of neurons expressing tdTomato, and the pre- and postsynaptic markers Bassoon and Homer. The image was composed of 4 × 7 tiled *z*-stacks acquired using a 60× 1.27 NA water-immersion objective and was stitched together to result in a ~380 × 650 × 50 µm^3^ volume in post-expansion space as shown **Fig. 4j**. High resolution information was also preserved as seen from the pre- and postsynaptic markers in **Fig. 4k**, revealing a ~150 nm separation (in pre-expansion dimensions) from the intensity profile as shown in **Fig. 4l**.

#### 3.4.2 SDCM + SMLM

Next, we explored the use of the DIY SDCM for the SMLM methods DNA-PAINT and STORM (stochastic optical reconstruction microscopy). For these SMLM measurements, we used Sector 2 of the disk (**Table 1**) which had been designed to use a slightly larger pinhole (*d* = 65 µm) for the longer wavelength emitting fluorophores (Cy3B and Cy5 or Alexa Fluor 647) and which used a closer spacing to help boost the fill factor to 0.056 for more efficient transmission of the illumination light with the tradeoff of reducing optical sectioning. The illumination light intensity was also boosted by using the high-intensity/small-area configurations of the illumination light path as indicated in **Fig. 1b-c**.

We first tested the SDCM via DNA-PAINT as it has been previously demonstrated using a commercial SDCM.[13] The sample was scanned initially by SDCM with a low labeling density conventional immunofluorescence stain of microtubules using the larger illumination area (80 × 80 µm^2^) at 488 nm as shown in **Fig. 5a**. Once a region of interest was identified for DNA-PAINT imaging, the beam was contracted to a 40 × 40 µm^2^ illumination area by switching the illumination telescope lenses (**Fig. 1b**) and the sample was imaged at 9.375 Hz using a 561 nm illumination power density of 135 W/cm^2^. Single-molecule binding events were observed with a characteristic dwell time of ~180 ms and an average of ~47,000 detected photons per localization. The resulting reconstructed image (green) was overlaid on the corresponding confocal *z*-section (magenta) in **Fig. 5b**. The characteristic hollow feature of immunolabeled microtubules by SMLM [27]was clearly visible throughout the entire image, even in areas containing multiple microtubules as shown in **Fig. 5c**. The peak-to-peak separation was fit to two Gaussian functions across 170 locations in the image resulting in an average separation of 41 ± 5 nm (**Fig. 5d-e**). The optical sectioning of SDCM reduces of out of focus background caused by the freely diffusing DNA-PAINT reporters.

**Figure 5.**
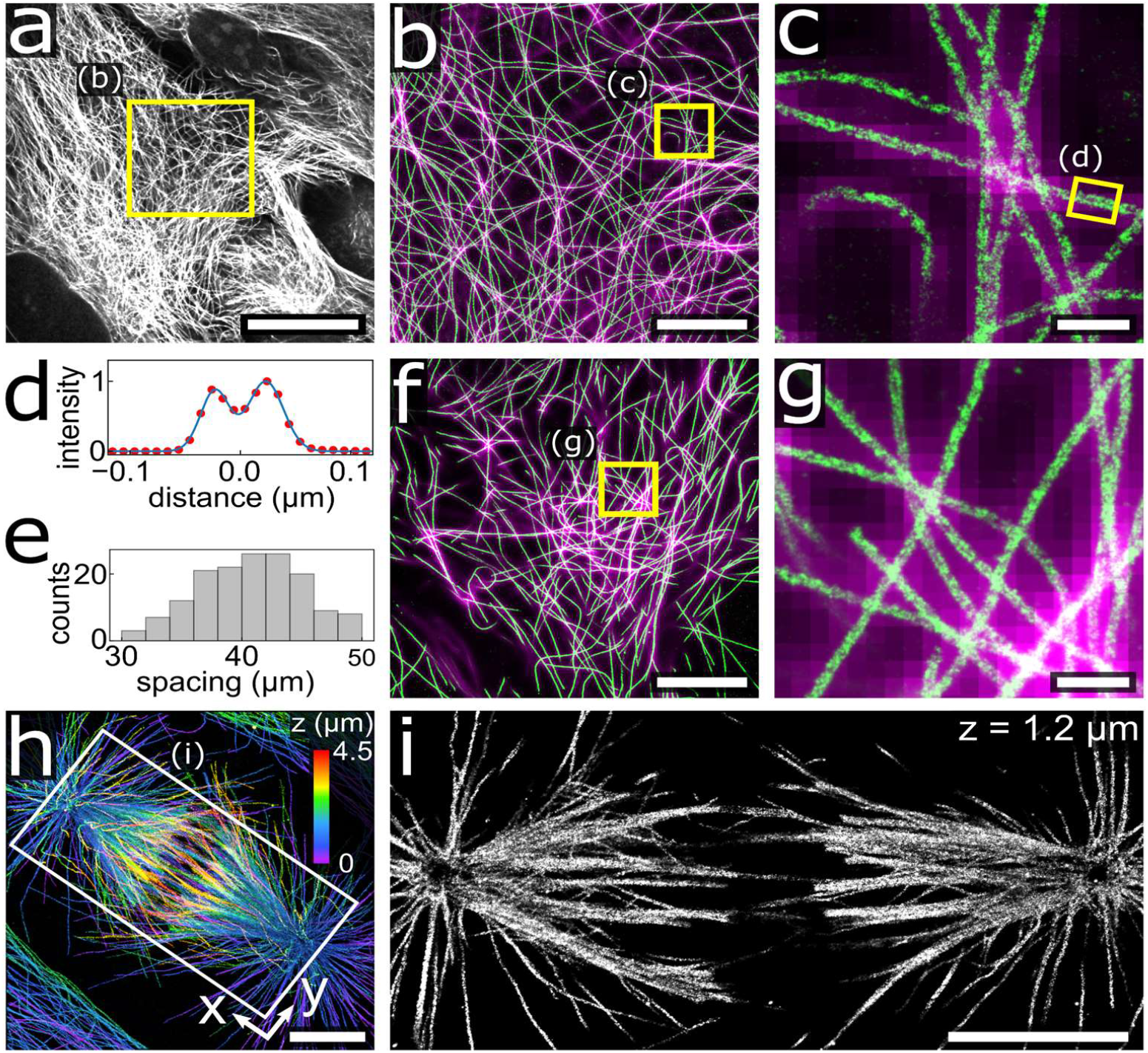
Demonstration of single molecule localization microscopy using SDC microscope. (a) Single confocal z-plane of immunostained microtubules of RPE-1 cells. (b) Overlay of DNA-PAINT image (green) acquired using an SDC, and the corresponding confocal z-plane (magenta) of the highlighted region in **a**. (c) Zoom in of the highlighted region in **b** showing microtubule hollow features. (d) Cross-sectional profile (dots) and dual Gaussian fit (solid) of boxed microtubule in **c**, showing a 43 nm peak-to-peak separation. (e) Histogram of peak-to-peak distances for 170 microtubules from panel **b**. (f) STORM image of immunostained microtubules in RPE-1 cells (green) collected through SDC, and overlay of the widefield epifluorescence image (magenta). (g) Zoom in of the region in **f**, showing ability to resolve microtubule hollow features. (h) Maximum intensity projection of 3D STORM image of immunostained microtubules from a 4.5 µm thick dividing PTK-1 cell with z-dimension position colorized according to the color scalebar. (i) Zoom in of a 50 nm z-section from the highlighted region in **h** at z = 1.2 µm. Scale bars, 20 µm (a), 5 µm (b,f,h,i), 500 nm (c,g).

While the application of commercial SDCM to DNA-PAINT has been previously shown, SMLM requiring photoswitching has had limited success using confocal microscopes and we aim to show its feasibility using DIY SDCM.[14,15] Using the high-intensity illumination configuration in **Fig. 1c**, we increased the 647 nm laser output power to 1500 mW, enabling us to achieve an excitation power density at the sample up to ~2 kW/cm^2^ on a prototype disk (70 µm pinholes and 350 µm inter-spiral spacing). With this illumination configuration, we were able to acquire good 15 Hz STORM movies on dense microtubule regions of PTK-1 cells which produced high-quality reconstructions (e.g., **Fig. 5f-g**). As with our SDCM DNA-PAINT measurements, the SDCM STORM reconstructed images prominently showed the characteristic hollow feature of the immunolabeled microtubules **Fig. 5g**.

Finally, we tested the sectioning ability of the SDCM for 3D STORM imaging by inserting a cylindrical lens into the detection path [19] and using a higher fill factor design (Sector 2) which enabled us to record movies at a frame rate of 37.5 Hz. We selected a mitotic PTK-1 cell for its increased thickness and densely labeled microtubules. We were able to acquire a series of STORM images at 200 nm steps across the full 4.5 µm volume encapsulating the mitotic spindle thickness totaling 1.2 million frames and ~16 million localizations. The resulting images were aligned and stitched as shown in the 3D maximum intensity projection in **Fig. 5h**. We observed large k-fiber bundles as well as individual microtubules in the interior of the spindle as shown in the single slice in **Fig. 5i** (see Fig. S4 for zoom-in views). For densely labeled samples, the optical sectioning provided by the SDCM is particularly helpful at removing background signal caused by out of focus molecules.

## 4. Discussion

The affordable DIY SDCM that we present here had strong performance in terms of spatial resolution, utility with various objective lenses, and application to a number of routine and challenging applications for biological imaging. Our own group has already used this DIY SDCM for imaging thin tissue sections and expanded cultured cells [40,41] although we have not previously described the DIY SDCM itself.

From PSF measurements, we observed the expected trends of improved lateral and axial resolution at shorter wavelengths with a constant (50 µm) pinhole size as shown in **Table 2**, with the exception that the 405 nm channel consistently had the worst axial resolution of the series. We speculate that this may be caused by poorly corrected axial aberrations in the objective lenses in this region of the spectrum. On the other hand, varying the pinhole size from 20 µm to 70 µm with excitation wavelengths 488 nm, 561 nm and 647 nm only led to small differences in resolution for the 100× 1.45 NA oil-immersion lens (**Table S3**). Instead, we found that the smallest pinhole sizes (~20-30 µm) exhibited substantially lower signal levels due to the compounded losses in excitation power and detected signal which may have obscured any resolution gains. While confocal microscopy is often stated to have improved lateral resolution by a factor of ~1.4 compared to epifluorescence microscopy, when the pinhole size is equal to or larger than the Airy diameter, as in this case, little to no resolution gain is to be expected. Instead, we saw much larger differences in resolution due to proper index matching for the 100× 1.45 NA oil-immersion lens via choice of immersion oil index of refraction in the range 1.500 to 1.530 or proper setting of the correction collar on the 60× 1.27 NA water-immersion lens (**Fig. S3**). [42]

The key design choice to use a SD without a microlens array enabled a build that was both simple and inexpensive but led to substantial losses of excitation laser power, for instance with only ~3.1% transmission (e.g., for sectors 3 and 5, **Table 1**) in comparison with >50% transmission for commercial designs. We mitigated these losses by taking two measures. First, we implemented a simple and inexpensive refractive beam shaper (**Fig. 1**) in order to reshape the input Gaussian beam into a flat beam with a square profile so that the available laser power would be distributed in the most efficient way possible. Second, we used a simple design in the illumination path that permitted 1:2 magnification or 2:1 demagnification of the square beam to enable a 16-fold variation in the laser illumination intensity. These measures were sufficient to achieve a range of power densities (**Table S3**) suitable for a variety of uses of the SDCM ranging from conventional imaging to single-molecule fluorescence microscopy (**Fig. 4, Fig. 5**).

While fixed immunostained cultured cells provided bright and robust specimens, in contrast, FISH specimens are limited in brightness as they generally label each mRNA with ~15-40 fluorophores depending on the probe set and hybridization efficiency. This low signal can be a challenge for conventional confocal microscopy, and SDCM may be better suited when optical sectioning is required. [43] Fortunately, while expanded specimens often appear dimmer by volumetric reduction of fluorophore density, in this case the mRNA puncta do not suffer from this effect because they generally remain smaller than the focal volume. The improved resolution allowed easier segmentation of individual mRNA, and the observation of some 3D structure in the pair of bright clusters in the nucleus ascribed to bursts of nascent mRNA as shown in **Fig. 4h-i**. Each mRNA appeared as a diffraction-limited spot, with ~20,000 mean detected photons per puncta for ~2,500 mRNAs analyzed. While SDCM may offer a powerful tool for mRNA FISH, good sample preparation was imperative. In the example here, the probe set contained 24 probes, giving the upper limit of 24 fluorophores labelling each transcript, and we found that ATTO 565 was particularly useful for the application.

A major benefit of SDCM is the acquisition speed as compared to conventional confocal microscopy. This is particularly advantageous for larger samples, such as cleared or expanded tissues, where the acquisition time scales volumetrically with sample size. The exposure time must be set with the sample in mind, taking into consideration factors such as the degree of labeling, photobleaching, available laser power and time resolution. For the expanded specimen, we were able to use exposure times down to 133 ms, corresponding to 10 complete revolutions of the disk. While the DIY SDC is still slower than the maximum frame rates achievable by commercial modules, it provides a reasonable 5-10× boost over conventional confocal microscopy, assuming a frame rate of ~1 Hz. The tiling presented in **Fig. 4j** was demonstrated as a proof of concept, but volumetric imaging would benefit from the further improvements of increasing the illumination area to utilize the full sCMOS chip, and from using a *z*-stage with a larger travel range to utilize the full ~170 µm working distance of the water-immersion objective.

The SMLM performance by SDCM was overall strong. As with a previous report using a commercial spinning disk system, we found that the spinning disk was helpful in rejecting out of focus light, although we also used a lower concentration reporter DNA oligonucleotide (50 pM) to help ensure the localizations were sufficiently sparse.[13] We were able to observe the characteristic hollowness of immunolabeled microtubules which appeared with an average separation of ~41 nm, slightly larger than the value typically reported (37 nm) likely due to the additional streptavidin linkage used immobilize the target DNA in our staining protocol.[27] SDCM enabled optical sectioning that was helpful when imaging thick and densely labeled specimens by STORM that more typically use TIRF or angled illumination to reduce out of focus background signal. With SDCM STORM, single molecules were observed with ~ 6,500 photons per localization and a characteristic on time of ~0.1 s. As a comparison, when we bypassed the disk and conducted STORM in epifluorescence at 200 Hz, localizations were detected with ~10,000 photons. Even so, the resolution standard of microtubule hollowness was clearly observed by SDCM STORM. The lower optimal frame rate of SDCM STORM was likely due to the fractional amount of area exposed (corresponding to the fill factor of the disk) as compared to epifluorescence.

## 5. Conclusions

At its core, SDCMs are built around a disk that spins, and the remaining microscope components differ little from a common laser-illuminated epifluorescence setup. Commercial SDCMs utilize a carefully registered microlens array to boost the excitation efficiency to compensate for low source power, but these come at significant additional cost and increase in complexity. Considering the current affordability of lasers that have sufficient power for SDs without a microlens, and the number of fluorescence setups already equipped with such lasers, a low-cost SD module using a glass photomask and readily available commercial components is well within reach of many labs. The SDCM can be easily designed to have multiple sectors that are optimized for objective lens choice or application (e.g., larger pinhole-pinhole distance to reduce pinhole-pinhole crosstalk for thicker samples or a larger fill factor for SMLM in thin specimens). The SDCM showed strong performance from the UV to NIR range, with different objective lenses, and with different modalities including conventional microscopy and super-resolution microscopy by specimen expansion, DNA-PAINT, or STORM. Our SDCM design is well-documented and can hopefully serve as encouragement to others with experience building optical instruments to create their own SDCMs to bring affordable high-performance capabilities to a wider group of researchers.

## Supporting information

Supplemental Codes

## Funding

National Institutes of Health (R01 MH115767).

## Acknowledgment

The authors acknowledge Brian Beliveau (Department of Genome Sciences, University of Washington, Seattle WA) for advice and DNA-PAINT reagents.

## Disclosures

The authors declare no conflicts of interest.

## Notes

### Competing Interest Statement

The authors have declared no competing interest.

