## Supplemental Document for "Versatile, do-it-yourself, low-cost spinning disk confocal microscope"

**Table S1. Component list for DIY spinning disk confocal microscope.**

| Component Name | Vendor | Catalog # | Quan. | Price ($) | Total ($) |
|---|---|---|---|---|---|
| **Lens** | | | | | |
| f=50 mm Achromatic Doublet | Thorlabs | AC254-050-A-ML | 1 | 106.05 | 106.05 |
| f=100 mm Achromatic Doublet | Thorlabs | AC254-100-A-ML | 1 | 106.05 | 106.05 |
| f=150 mm Achromatic Doublet | Thorlabs | AC254-150-A-ML | 3 | 106.05 | 318.15 |
| **Optomechanics** | | | | | |
| 30 mm Cage Cube with Dichroic Filter Mount | Thorlabs | CM1-DCH | 1 | 175.31 | 175.31 |
| Kinematic Prism Mount | Thorlabs | KM100PM | 1 | 80.07 | 80.07 |
| Mounted Standard Iris | Thorlabs | ID25 | 1 | 60.87 | 60.87 |
| Precision Optical Rail | Newport | PRL-6 | 1 | 165 | 165 |
| Rail Carrier PRL Series | Newport | PRC-3 | 1 | 107 | 107 |
| **Filters, Dichroics and Wheel** | | | | | |
| Laser polychroic mirror | Chroma | zt405/488/561/647/752rpc-UF2 | 1 | 975 | 975 |
| Emission Filter ET445/58m | Chroma | ET445/58m | 1 | 325 | 325 |
| Emission Filter ET525/50m | Chroma | ET525/50m | 1 | 325 | 325 |
| Emission Filter ET605/70m | Chroma | ET605/70m | 1 | 325 | 325 |
| Emission Filter ET700/75 | Chroma | ET700/75 | 1 | 325 | 325 |
| Emission Filter ET800/60 | Chroma | ET800/60 | 1 | 325 | 325 |
| Motorized Filter Wheels | Thorlabs | FW102C | 1 | 1211.97 | 1211.97 |
| **Mirrors** | | | | | |
| Broadband Dielectric Mirror | Thorlabs | BB1-E02 | 5 | 77.35 | 386.75 |
| **Beam Shaper** | | | | | |
| Topag top hat beam shaper, 250mm WD 4x4mm top hat 400-700nm AR/AR | CourierTronics | GTH-5-25-4 | 1 | 668 | 668 |
| Topag positioner for GTH-5-25-4 | CourierTronics | HSF01 | 1 | 176 | 176 |
| **Photomask for Spinning Disk** | | | | | |
| 5X5X.090 Quartz ND5 10um Resolution, cut and center hole, both sides BBAR | Front Range Photomask | NA | 1 | 974 | 974 |
| **Fasteners and Miscellaneous** | | | | | |
| Screw 2-56 Thread Size, 3/8" | McMaster | 91253A079 | 1 | 6.09 | 6.09 |
| O-Ring Dash Number 016 | McMaster | 9452K6 | 1 | 4.46 | 4.46 |
| O-Ring Dash Number 022 | McMaster | 9452K76 | 1 | 5.93 | 5.93 |
| Yeeco 5-12V BLDC motor controller | Amazon | B07BQYYDPH | 1 | 12.99 | 12.99 |

| | | | | | |
|---|---|---|---|---|---|
| | | | | **Grand Total** | **7164.69** |

| **Alternative or Optional Components** | | | | | |
|---|---|---|---|---|---|
| 5x5x.090 LRC Chrome 10um Resolution Mask, cut and center hole | Front Range Photomask | NA | 1 | 392 | 392 |
| Laser polychroic mirror | Chroma | ZT405/488/561/647rpc | 1 | 550 | 550 |
| Bandpass Emission Filter | Chroma | ET605/70m-2p | 1 | 475 | 475 |
| Bandpass Emission Filter | Chroma | ET525/50m-2p | 1 | 475 | 475 |
| Externally SM1-Threaded End Cap | Thorlabs | SM1PL | 2 | 15.48 | 30.96 |
| Externally SM1-Threaded Plug | Thorlabs | SM1CP2 | 2 | 18.83 | 37.66 |

**Table S2. Summary of sample preparation and imaging conditions.**

| **Figure** | **Specimen** | **1° Ab** (1-2 µg/mL) **or FISH** | **2° Ab** (2-5 µg/mL or as indicated) **or FISH** | **Additional Stains** | **Imaging Cocktail** | **Objective lens** | **Disk Sector** | **Illumination & Area** | **Intensity, exposures, Z-step size** |
|---|---|---|---|---|---|---|---|---|---|
| **3 a-d** | Beads (fluoresbrite YG 0.1 µm) | | | | | 100× 1.45 NA oil (n = 1.512) | 3 | 2× expand<br>80×80 $\mu m^2$ | 488 nm: 18 $W/cm^2$<br>133 ms exposures |
| **3 f-g** | 1M fluorescein solution | | | | | 100× 1.45 NA oil (n = 1.512) | prototype disk<br>*d* = 50 µm<br>*s* = 250 µm<br>or<br>*s* = 500 µm | | |
| **4 a-f** | BS-C-1 cells, extracted then PFA/GA fixed | Rat x tTub<br>Rb x dTub<br>Ms x Vim | D x Rat AF750 (3.5 d/p)<br>D x Rb AF568 (7.5 d/p)<br>D x Ms AT488 (6 d/p) | Phalloidin AF647 0.6 µM, Hoechst 2 µg/mL in PBS | Glox + 1 mM trolox | 60× 1.27 NA water | 3 | 2× expand<br>130×130 $\mu m^2$ | 405: 2 $W/cm^2$<br>488: 7 $W/cm^2$<br>561: 6 $W/cm^2$<br>647: 4 $W/cm^2$<br>750: 7 $W/cm^2$<br>266 ms exposures<br>250 nm z-steps |
| **4 g-i** | RPE-1 cells, PFA fixed | GAPDH probe set, 150 nM, overnight 37 °C | Sequence P5 AT565 reporter oligo, 20 nM in 2× SSC | TO-PRO-3 1 µM | 0.02× SSC<br>(3× expansion) | 60× 1.27 NA water | 5 | 2× expand<br>130×130 $\mu m^2$ | 561: 32 W/cm2<br>647: 2 W/cm2<br>266 ms exposures<br>250 nm z-steps |
| **4 j-k** | Perfusion PFA fixed mouse brain,<br>100 µm slice | Rb x Homer<br>Ms x Bassoon<br>Rat x mCherry | D x Rb AF488 (8.5 d/p)<br>D x Ms AT647N (3.5 d/p)<br>D x Rat AT565 (3.8 d/p) | | water (4× expansion) | 60× 1.27 NA water | 5 | 2× expand<br>130×130 $\mu m^2$ | 488: 5 $W/cm^2$<br>561: 10 $W/cm^2$<br>647: 7 $W/cm^2$<br>266 ms exposures<br>500 nm-z steps |
| **5 a-c** | RPE-1 cells, extracted then PFA/GA fixed | Rat x Tub | D x Rat Biotin (2 µg/mL)<br>D x Rat AT488 (7 d/p, 0.2 µg/mL) | streptavidin 5 µg/mL, then sequence P1C biotin 500 nM | PAINT imaging buffer<br>Sequence P1 Cy3B 0.05 nM | 100× 1.45 NA oil | 2 | None<br>40×40 $\mu m^2$ | 488: 10 $W/cm^2$<br>561: 135 $W/cm^2$<br>106 ms exposures |
| **5 f-g** | B-SC-1 cells, extracted, then PFA/GA fixed | Rat x Tub | D x Rat AF647 (4.6 d/p) | | Glox + 143 mM βME | 100× 1.45 NA oil | prototype disk<br>*d* = 70 µm<br>*s* = 350 µm | 2× contract<br>20×20 $\mu m^2$ | 647: 1 $kW/cm^2$<br>66 ms exposures |
| **5 h,i** | PtK-1 cells, extracted, then PFA/GA fixed | Rat x Tub<br>Ms x Tub | D x Rat AF647 (4.6 d/p)<br>D x Ms AF647 (5.3 d/p) | | Glox + 143 mM βME | 100× 1.45 NA oil | 2 | 2× contract<br>20×20 $\mu m^2$ | 647: 1.2 $kW/cm^2$<br>26 ms exposures |
| **S3, Table 2** | Beads (Tetraspeck 0.1 µm or 0.2 µm) | | | | Optical cement<br>Water<br>Air<br>Air | 100× 1.45 NA oil<br>60× 1.27 NA water<br>100× 0.9 NA air<br>20× 0.45 NA air | Various sectors | 2× expand<br>80×80 $\mu m^2$ | all: ~20 $W/cm^2$<br>266 ms exposures |
| **S4, Table 2** | BS-C-1 cells, extracted, then PFA/GA fixed | Rat x Tub | D x Rat AF405 (2 d/p)<br>D x Rat AT488 (7 d/p)<br>D x Rat AF568 (3.4 d/p)<br>D x Rat AF647 (4.6 d/p)<br>D x Rat AF750 (3.5 d/p) | | Glox + 1 mM trolox<br>or<br>Glox + 1 mM trolox + 60% sucrose for oil imaging | 100× 1.45 NA oil<br>60x 1.27 NA water<br>100x 0.9 NA air<br>20x 0.45 NA air | Various sectors | 2× expand<br>80×80 $\mu m^2$ or 130×130 $\mu m^2$ | all: ~10 $W/cm^2$<br>266 ms exposures |

**Abbreviations:** 1° Ab = primary antibody; 2° Ab = secondary antibody; AF = Alexa Fluor; AT = ATTO-TEC; *d* = disk pinhole diameter; d/p = average number of dye molecules per protein; D x Rat = donkey anti-rat IgG; D x Rb = donkey anti-rabbit IgG; D x Ms = donkey anti-mouse IgG; GA = glutaraldehyde; Glox = glucose oxidase and catalase mixture; NA = numerical aperture; PBS = phosphate buffered saline; PFA = paraformaldehyde; *s* = inter-spiral spacing; SSC = saline sodium citrate; tTub = tyrosinated tubulin; dTub = detyrosinated tubulin; Vim = vimentin.

**Notes:** Illumination intensities are reported as average intensity across the field of view.

**Table S3. DNA sequences for GAPDH mRNA FISH and DNA PAINT.**

| GAPDH Probes | Sequence |
|---|---|
| GAPDH_P01 | GGGTGGAATCATATTGGAACATGTAAACCATGTTAATGATACGGCGACCACCGA |
| GAPDH_P02 | AGCCAAATTCGTTGTCATACCAGGAAATGAGCTTAATGATACGGCGACCACCGA |
| GAPDH_P03 | TGGTGATGGGATTTCCATTGATGACAAGCTTCTTAATGATACGGCGACCACCGA |
| GAPDH_P04 | TTTACCAGAGTTAAAAGCAGCCCTGGTGACTTAATGATACGGCGACCACCGA |
| GAPDH_P05 | TTTATTGATGGTACATGACAAGGTGCGGCTCCTTAATGATACGGCGACCACCGA |
| GAPDH_P06 | CAAAAGAAGATGCGGCTGACTGTCGAATTAATGATACGGCGACCACCGA |
| GAPDH_P07 | TTCAAGGGGTCTACATGGCAACTGTGATTAATGATACGGCGACCACCGA |
| GAPDH_P08 | TTCTCATGGTTCACACCCATGACGAACATGGTTAATGATACGGCGACCACCGA |
| GAPDH_P09 | GGCATTGCTGATGATCTTGAGGCTGTTGTCTTAATGATACGGCGACCACCGA |
| GAPDH_P10 | ACGCCTGCTTCACCACCTTCTTGATGTCATCATTAATGATACGGCGACCACCGA |
| GAPDH_P11 | TCTCAGCCTTGACGGTGCCATGGAATTTGTTAATGATACGGCGACCACCGA |
| GAPDH_P12 | ACTTGATTTTGGAGGGATCTCGCTCCTGGTTAATGATACGGCGACCACCGA |
| GAPDH_P13 | GGACTGTGGTCATGAGTCCTTCCACGATATTAATGATACGGCGACCACCGA |
| GAPDH_P14 | CAAAGTGGTCGTTGAGGGCAATGCCATTAATGATACGGCGACCACCGA |
| GAPDH_P15 | AGTTGTCATGGATGACCTTGGCCAGGTTAATGATACGGCGACCACCGA |
| GAPDH_P16 | CAGTAGAGGCAGGGATGATGTTCTGGTTAATGATACGGCGACCACCGA |
| GAPDH_P17 | TCGCTGTTGAAGTCAGAGGAGACCACTTAATGATACGGCGACCACCGA |
| GAPDH_P18 | GACCAAATCCGTTGACTCCGACCTTCACCTTAATGATACGGCGACCACCGA |
| GAPDH_P19 | TTCTCCATGGTGGTGAAGACGCCAGTGTTAATGATACGGCGACCACCGA |
| GAPDH_P20 | GTCTTACTCCTTGGAGGCCATGTGGGTTAATGATACGGCGACCACCGA |
| GAPDH_P21 | CCACAGTCTTCTGGGTGGCAGTGATGTTAATGATACGGCGACCACCGA |
| GAPDH_P22 | GTTCAGCTCAGGGATGACCTTGCCCATTAATGATACGGCGACCACCGA |
| GAPDH_P23 | CAGGTCAGGTCCACCACTGACACGTTTTAATGATACGGCGACCACCGA |
| GAPDH_P24 | CTCAGTGTAGCCCAGGATGCCCTTGATTAATGATACGGCGACCACCGA |
| **GAPDH Reporter** | ATTO565-CTGTAGCAGTTCGGTGGTCGCCGTATCATT |
| **DNA PAINT** | |
| P1C (docking) | Biotin-TTTTTATACATCTA |
| P1 (imaging) | CTAGATGTAT-Cy3B |

**Table S4. Summary of Point Spread Functions for different disk and objective combinations.**

| Pinhole Dia (µm) | $\lambda_{ex}$ | X-FWHM (µm) | stdX-FWHM | Z-FWHM (µm) | stdZ-FWHM | Sample |
|---|---|---|---|---|---|---|
| **20× 0.45 NA Air** | | | | | | |
| 35 | 488 | 0.80 | 0.06 | 4.89 | 0.19 | TS200 |
| 35 | 561 | 0.84 | 0.05 | 4.95 | 0.11 | " |
| 35 | 647 | 0.85 | 0.03 | 5.38 | 0.07 | " |
| 50 | 488 | 0.75 | 0.03 | 4.72 | 0.20 | " |
| 50 | 561 | 0.80 | 0.02 | 5.32 | 0.21 | " |
| 50 | 647 | 0.88 | 0.01 | 6.23 | 0.17 | " |
| **100× 0.9 NA Air** | | | | | | |
| 35 | 488 | 0.298 | 0.008 | 0.69 | 0.01 | TS100 |
| 35 | 561 | 0.325 | 0.009 | 0.78 | 0.02 | " |
| 35 | 647 | 0.370 | 0.008 | 0.89 | 0.04 | " |
| 50 | 488 | 0.317 | 0.010 | 0.74 | 0.01 | " |
| 50 | 561 | 0.334 | 0.007 | 0.81 | 0.01 | " |
| 50 | 647 | 0.386 | 0.008 | 0.93 | 0.02 | " |
| **100× 1.27 NA Water** | | | | | | |
| 35 | 488 | 0.32 | 0.01 | 0.77 | 0.04 | TS100 |
| 35 | 561 | 0.34 | 0.02 | 0.74 | 0.03 | " |
| 35 | 647 | 0.337 | 0.008 | 0.72 | 0.01 | " |
| 50 | 405 | 0.30 | 0.05 | 1.37 | 0.08 | " |
| 50 | 488 | 0.308 | 0.008 | 0.82 | 0.04 | " |
| 50 | 561 | 0.33 | 0.01 | 0.83 | 0.04 | " |
| 50 | 647 | 0.353 | 0.005 | 0.85 | 0.02 | " |
| 50 | 750 | 0.44 | 0.03 | 1.04 | 0.05 | " |
| **100× 1.45 NA Oil** | | | | | | |
| 20* | 488 | 0.24 | 0.02 | 0.40 | 0.01 | TS100 |
| 20* | 561 | 0.25 | 0.02 | 0.40 | 0.02 | " |
| 20* | 647 | 0.25 | 0.02 | 0.47 | 0.04 | " |
| 30* | 488 | 0.22 | 0.01 | 0.42 | 0.01 | " |
| 30* | 561 | 0.23 | 0.01 | 0.41 | 0.01 | " |
| 30* | 647 | 0.250 | 0.008 | 0.48 | 0.02 | " |
| 35 | 488 | 0.207 | 0.005 | 0.42 | 0.01 | " |
| 35 | 561 | 0.225 | 0.005 | 0.42 | 0.01 | " |
| 35 | 647 | 0.270 | 0.006 | 0.49 | 0.02 | " |
| 40* | 488 | 0.224 | 0.007 | 0.45 | 0.01 | " |
| 40* | 561 | 0.236 | 0.009 | 0.45 | 0.01 | " |
| 40* | 647 | 0.260 | 0.007 | 0.51 | 0.02 | " |
| 50 | 405 | 0.22 | 0.01 | 0.96 | 0.16 | MT |
| 50 | 488 | 0.222 | 0.004 | 0.49 | 0.02 | TS100 |
| 50 | 561 | 0.237 | 0.003 | 0.50 | 0.02 | " |
| 50 | 647 | 0.273 | 0.003 | 0.56 | 0.02 | " |
| 50 | 750 | 0.35 | 0.02 | 0.61 | 0.03 | MT |
| 70* | 488 | 0.229 | 0.001 | 0.51 | 0.01 | " |
| 70* | 561 | 0.244 | 0.005 | 0.52 | 0.01 | " |
| 70* | 647 | 0.276 | 0.004 | 0.60 | 0.01 | " |

**Notes:**

1) X-FWHM and stdX-FWHM refer to the average lateral full-width at half maximum from Gaussian fits and the standard deviation from many such fits. Z-FWHM and stdZ-FWHM refer to the average axial full-width at half maximum from Gaussian fits and the standard deviation from many such fits

2) Point spread function widths were determined from either 100 nm Tetraspeck beads (TS100), 200 nm Tetraspeck beads (TS200), or microtubule samples (MT).

3) * denotes a disk without an anti-reflection coating.

4) The inter-spiral spacing for all sectors was equal to 5x the pinhole diameter.

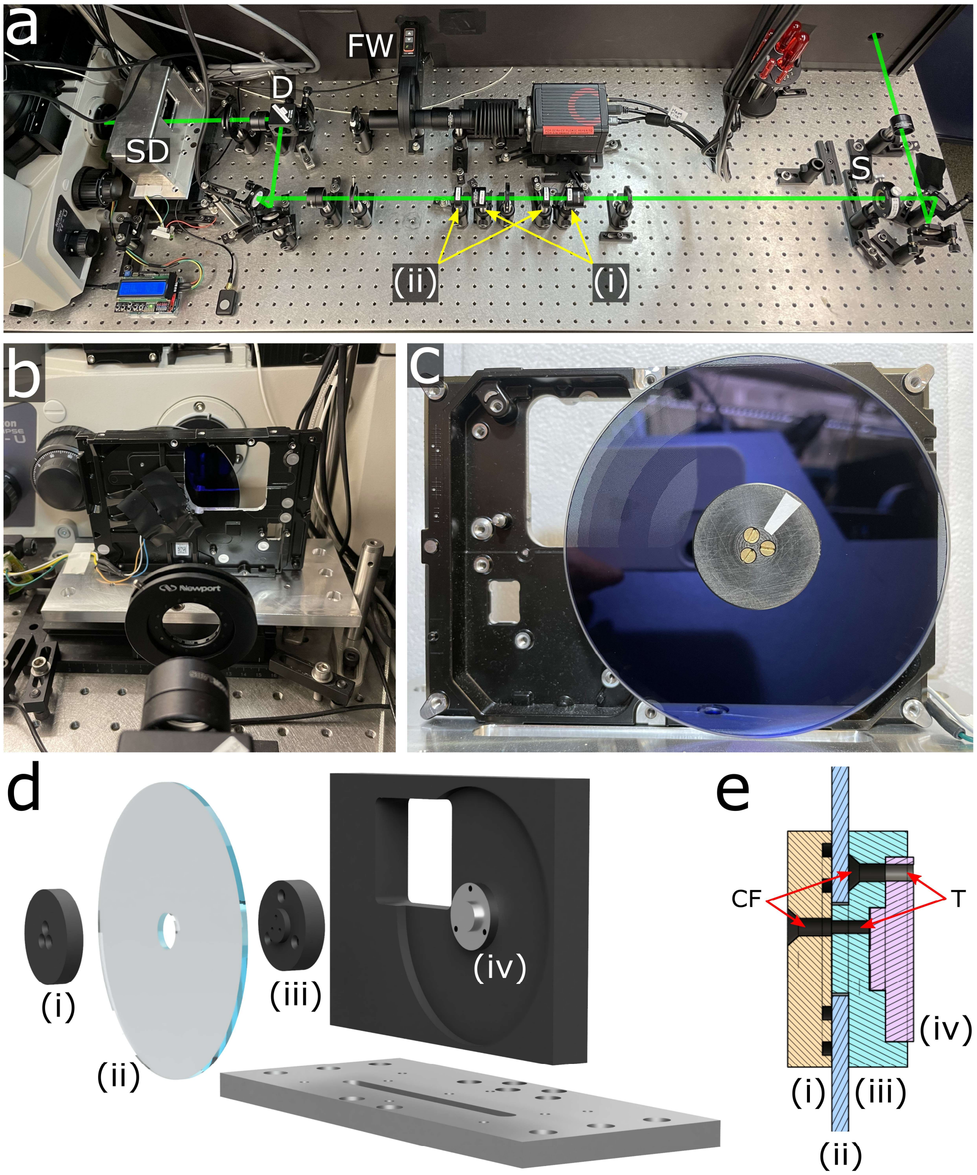

***Figure S1.*** *DIY SDCM setup and schematic diagrams. (a) Photograph of DIY SDCM with selected components highlighted: Spinning disk (SD) in its protective housing, dichroic (D), filter wheel (FW) and square shaper (S). The excitation laser path is highlighted in green, with 2× expanding telescope lenses (i) for general illumination, or 2× demagnifying telescope lenses for high power density (STORM illumination) by reversing the lens order in the marked optical mounts (ii). Alternatively, the telescope lenses can be omitted and the shaper repositioned for no magnification (DNA-PAINT illumination). (b) Photograph of spinning disk with housing removed. The commercial hard drive motor-base assembly is modified to include a machined window for the light path. (c) Photograph of the spinning disk showing the Delrin adapter and white tape mark used to trigger the tachometer circuit. (d) Schematic diagram and (e) cross-sectional view of spinning disk components and assembly. Marked components in (d) and (e) include (i) Delrin clamping plate, (ii) glass photomask, (iii) Delrin motor adapter, and (iv) hard drive spindle motor. Clearance and threaded holes are denoted CF and T, respectively. Note the O-ring grooves in the clamping plate ensure a strong grasp on the glass disk (see* ***Table S1*** *for part numbers). Refer to Code 3 for CAD files.*

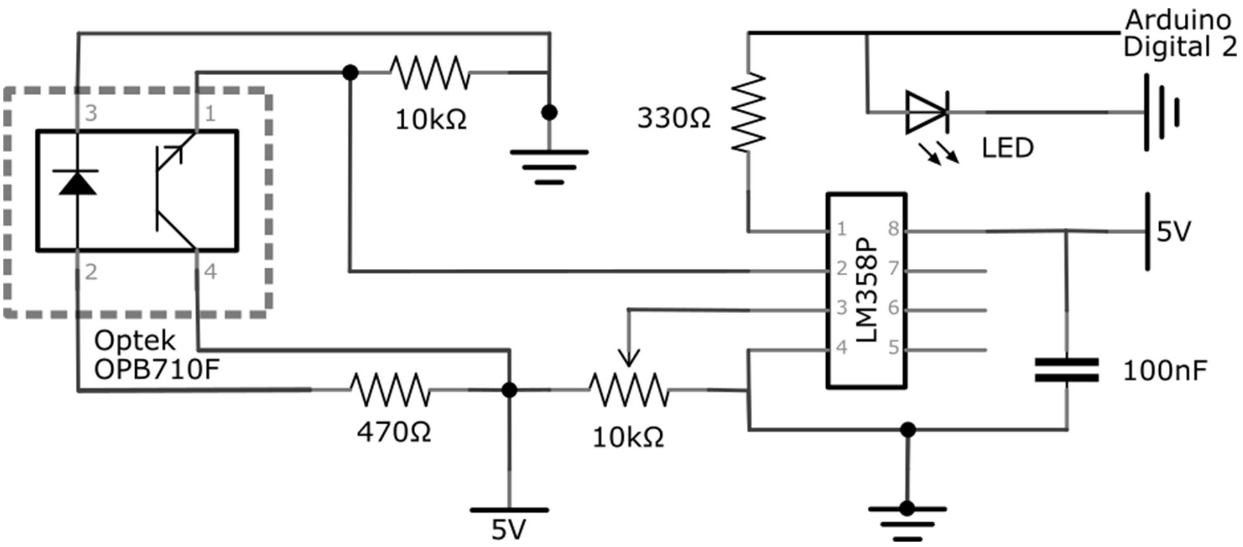

***Figure S2.*** *Circuit schematic of spinning disk tachometer. A reflective phototransistor photodiode pair (Optek, OPB710F, 910 nm) was fitted into the spinning disk housing above the Delrin clamping plate, and a small strip of absorptive tape placed applied to provide a reflective signal difference. The detection threshold of the operational amplifier (LM358P) was adjusted using the potentiometer, and a LED provided a visual readout. The digital output signal of the circuit was wired to a microcontroller (Arduino Uno) which counted the pulses in a predetermined time window, and the rotation rate was displayed on an LCD shield (DFRobot, DFR0009). The reflective sensor (dashed box) was connected on long flexible wires for ease of placement. Practically, we found it was unnecessary to trigger the camera from the disk rotation signal, instead setting an exposure time approximately equal to an integer number of disk rotations was sufficient to avoid any streaking artifacts. Refer to Code 4 for corresponding Arduino code.*

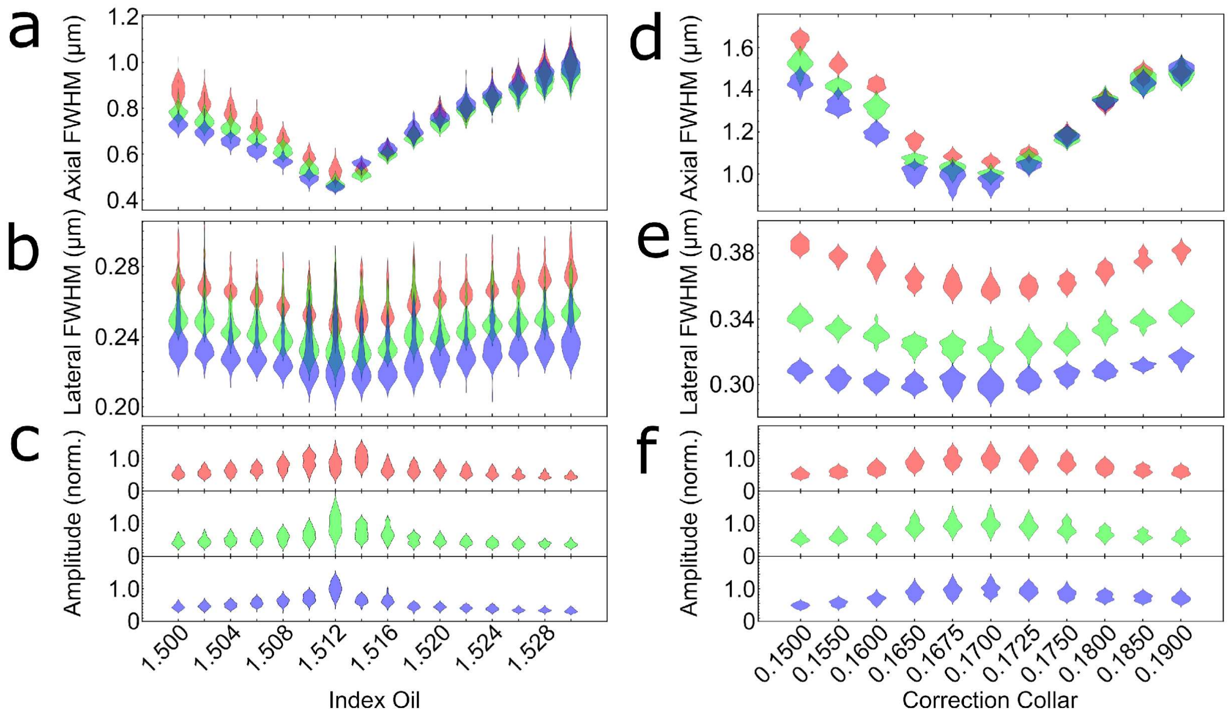

***Figure S3.*** *PSF axial FWHM, lateral FWHM, and maximum amplitude by varying index matching oil of a 100× 1.45NA oil immersion objective (a-c), or by varying the correction collar of a 60× 1.27NA water immersion objective (d-f), respectively. The analysis was performed on 100 nm Tetraspeck beads excited at 488 nm (blue), 561 nm (green) and 647 nm (red) using d = 50 µm and s = 250 µm. For the oil lens (a-c), the beads were adsorbed to a coverglass and embedded in optical cement (Nordland NOA60), and >90 beads were analyzed for each index oil. For the water lens (d-f), the beads were adsorbed to a coverglass, covered with water, and >60 beads were imaged for each collar position.*

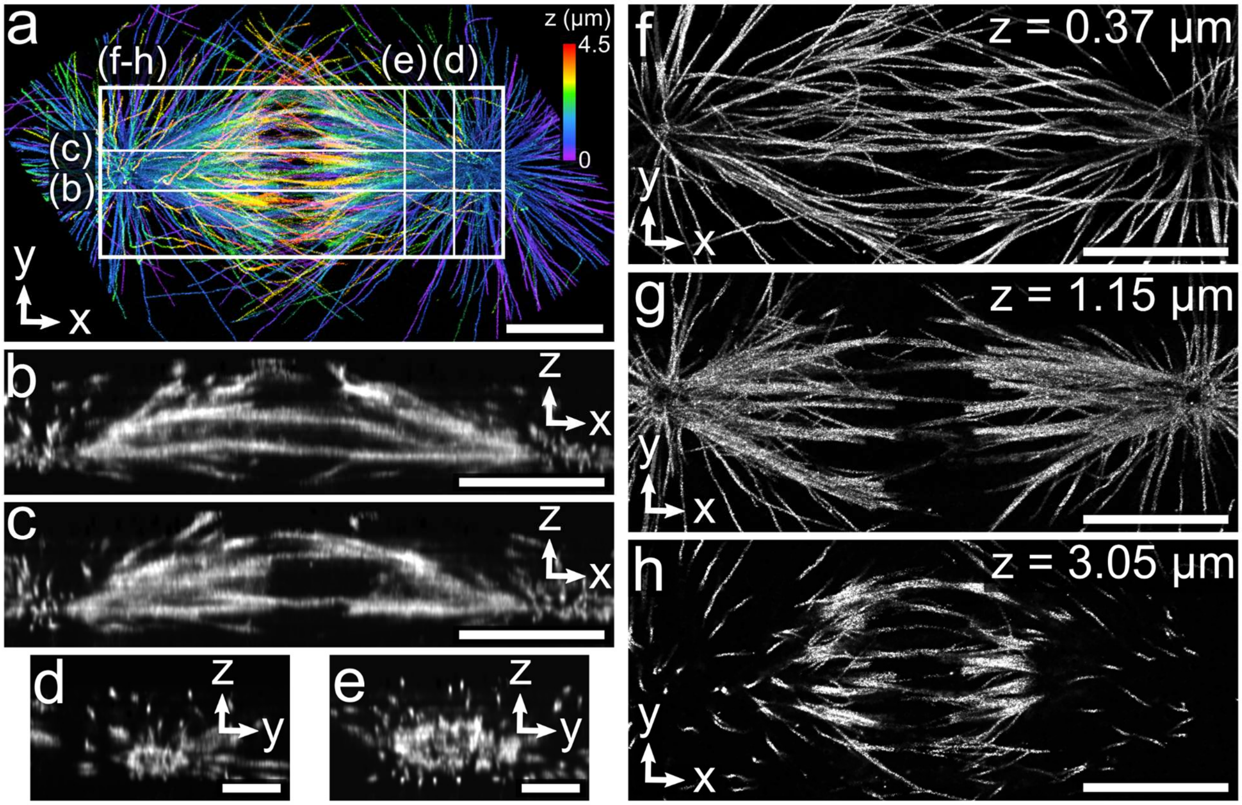

***Figure S4.*** *Single-molecule localization microscopy of a dividing PTK-1 cell using SDC microscope. (a) Maximum intensity projection of 3D STORM image of immunostained microtubules from a 4.5 µm thick dividing PTK-1 cell with z-dimension position colorized according to the color scale bar. (b-e) Cross-section corresponding to the planes denoted in **a**. (f) Maximum intensity projection of a 0.75 µm thick image centered at z=0.37 µm and (g,h) 50 nm thick z-sections corresponding to the z-height denoted in the figure for the area highlighted in **a**. Scale bars 5 µm (a-c, f-h), 2 µm (d,e).*

**Supplemental Note 1**

The alignment process is analogous to that of an epifluorescence microscope for most steps, however, in this design, the dichroic is placed external to the microscope chassis, and an empty cube is used in microscope turret. Once the microscope is set up for epifluorescence imaging in this way, two unique steps to the SDCM are described as follows. First, the spinning disk pinholes must be carefully placed at the image plane of the microscope. The positions of $L_4$, $L_5$ and the camera can be set without the disk in the emission path. Next, the disk is inserted near the image plane using white light illumination provided by the microscope condenser. The position of the disk at the image plane is set by carefully sliding it along the optical axis until the stationary pinholes are in sharp focus on the camera to determine the image plane. Observing the pinholes in sharp focus may be facilitated using larger pinholes (>50 µm) if available. Second, the iris between $L_4$ and $L_5$ can be stopped down to lower background, but not appreciably reduce fluorescence signal. In addition, the spinning disk can then be rotated slightly off perpendicular to the optical axis such that the reflected excitation beam is rejected by the iris. Of final note, we observed that it was best to face the reflective metal side of the spinning disk toward the camera to reduce any autofluorescence background generated by the disk substrate.
