## Supplemental Codes for "Versatile, do-it-yourself, low-cost spinning disk confocal microscope": SD_Tachometer.ino

```
#include <LiquidCrystal.h>

LiquidCrystal lcd(8, 9, 4, 5, 6, 7);        // select the pins used on the LCD panel

volatile float REVshort = 0; // VOLATILE DATA TYPE TO STORE REVOLUTIONS
volatile float REVlong = 0; // VOLATILE DATA TYPE TO STORE REVOLUTIONS

unsigned int shortinterval = 1000; // short term measurement interval in ms
unsigned int longinterval = 10000; // long term measurement interval in ms

float prevtimeshort = 0; // previous time of short interval measurement
float prevtimelong = 0; // previous time of long interval measurement

float hzs = 0.; //short inteval hz value
float hzl = 0.; //long inteval hz value

void setup(){
  lcd.begin(16, 2);                       // start lcd
  attachInterrupt(digitalPinToInterrupt(2), RPMCount, RISING); // Interrupt on Pin2
  lcd.print("SDC TACH"); // STARTUP TEXT
  delay(2000);
  lcd.clear();
  prevtimeshort = millis();
  prevtimelong = millis();
}

void loop()
{
 float currtime = millis(); // current time
 float timedifshort = currtime - prevtimeshort; // time between last measurement
 float timediflong = currtime - prevtimelong; // time between last measurement

 if(timediflong > longinterval) // measure after long interval
 {
    hzl = REVlong/(timediflong/1000.); // measured HRZ
    prevtimelong = currtime;
    REVlong = 0.;
 }
 
 if(timedifshort > shortinterval) // measure after shot interval
 {
    hzs = REVshort/(timedifshort/1000.); // measured HZ
    prevtimeshort = currtime;
    REVshort = 0.;
    
 if(hzs > 1)
 {
    lcd.clear();
    lcd.setCursor(0,0);
    lcd.print("hz(1s) : ");
    lcd.print(hzs);
    
    lcd.setCursor(0,1);
    lcd.print("hz(10s): ");
    lcd.print(hzl);
 }
 else
 {
    lcd.clear();
    lcd.print("disk spinning?");
 }
 }
}
 
void RPMCount() // EVERYTIME WHEN THE SENSOR GOES FROM LOW TO HIGH , THIS FUNCTION WILL BE INVOKED
{
 REVshort++; // INCREASE REVOLUTIONS
 REVlong++; // INCREASE REVOLUTIONS
}
```
